# Wasting food is disgusting: Evidence from behavioral and neuroimaging study of moral judgment of food-wasting behavior

**DOI:** 10.1101/750299

**Authors:** Michalina Marczak, Artur Marchewka, Marek Wypych, Michał Misiak, Dawid Droździel, Piotr Sorokowski, Agnieszka Sorokowska

**Author notes:** Correspondence concerning this article should be addressed to Agnieszka Sorokowska, University of Wroclaw, Institute of Psychology, ul. Dawida 1, 50-527 Wroclaw, Poland Contact.

## Abstract

Food-wasting has a profound negative social and environmental impact. Acknowledging that referring to moral judgment can motivate behavior change, the present study aimed to determine moral intuitions underlying the perception of food-wasting behavior. We developed a set of affective standardized scenarios and we used them to collect behavioral and neuroimaging data. In the main study, 50 participants made moral judgments regarding food-wasting, disgusting, harmful, dishonest, or neutral behaviors presented in these scenarios. We found that wasting food was considered morally wrong and it was associated with moral disgust. Neuroimaging data revealed that food-wasting stimuli elicited an increased activity in structures associated with moral judgment, as well as in regions involved in the processing of moral, but also physical disgust. We discuss our results in the context of the evolutionary significance of food that might have led to seeing food-wasting as a moral transgression.

---

It is estimated that approximately one-third of all produced food is lost during production and consumption (Gustavsson, Cederberg, & Sonesson, 2011). This problem has profound social and environmental implications. According to the World Economic Forum (2016), food-wasting practices interfere with efforts to establish global food security and may result in future food crises. In addition, food-wasting is associated with food overproduction, which contributes to deforestation and decreased biodiversity through rapidly increasing logging to create pastures and croplands (Houghton, 2012). Moreover, decomposing food is a source of methane which is a much more potent greenhouse gas than carbon dioxide (Melikoglu, Lin, & Webb, 2013). Thus, addressing the problem of food waste from various perspectives is necessary to reduce human malnutrition and biodegradation (Erb et al., 2016; Willett et al., 2019), both of which are among the largest current global risks associated with food-wasting (World Economic Forum, 2016).

Food is lost or wasted in every step of the supply chain. However, in developed countries, the largest waste contributors are direct consumers (Parfitt, Barthel, & Macnaughton, 2010). For example, in the European Union, 89 million tons of food is wasted annually, which is the equivalent to 173 kilograms of food waste per person. Over 50% of this food waste is produced in consumer households (Stenmarck et al., 2016). At the same time, it is estimated that 80% of food waste is avoidable, i.e., it is edible food which is not consumed (Vanham, Bouraoui, Leip, Grizzetti, & Bidoglio, 2015). Studies conducted in the United Kingdom (WRAP, 2007) and Australia (Hamilton, Denniss, & Baker, 2005) on the demographics of food waste show that young people waste more food than older people and that retired families with fewer members waste less food. Education also influences food waste: a lower level of education corresponds to a lower level of food waste (Monier et al., 2010).

However, despite the prevalence of food-wasting behaviors, only recently has this problem been approached from a psychological perspective (Farr-Wharton, Choi, & Foth, 2014; Farr-Wharton, Foth, & Choi, 2014; Graham-Rowe, Jessop, & Sparks, 2015; Kallbekken & Sælen, 2013; Porpino, Wansink, & Parente, 2016; Schanes, Dobernig, & Gözet, 2018; Stancu, Haugaard, & Lähteenmäki, 2016). It has been found, for example, that the largest concerns regarding food-wasting in industrialized societies are of a financial nature (Graham-Rowe et al., 2015; Neff, Spiker, & Truant, 2015; Quested, Marsh, Stunell, & Parry, 2013). Some researchers have begun to view food-wasting as a moral problem (Graham-Rowe et al., 2015; Misiak, Butovskaya, & Sorokowski, 2018). In a qualitative study by Graham-Rowe, Jessop, and Sparks (2014), while respondents were motivated to reduce household food waste principally to save money, moral concerns were indicated as the second-most important motivation. People felt that it was the “right thing to do” to reduce household food waste. When asked why they believed that food-wasting was wrong, participants provided several reasons, e.g., respect for tradition or concern for the environment and future generations.

People in traditional, small-scale societies have been found to associate wasting food with harming others. Misiak and colleagues (2018) hypothesized that moral judgments about food-wasting may serve as an adaptation to harsh ecologies and that populations with higher levels of food insecurity may develop moral norms which prevent people from wasting food. These authors report that the Maasai, who deal with food shortages on a daily basis, tend to compare food-wasting to physical harm (e.g. food-wasting is worse than killing, because killing harms only one person and wasting food may cause more deaths).

Nevertheless, a simple statement that wasting food is immoral is not exhaustive. Moral judgment is a complex, non-unitary phenomenon that can be studied in multiple ways, including in a neuropsychological perspective (Awad et al., 2018; Bloom, 2010; Crockett, Siegel, Kurth-Nelson, Dayan, & Dolan, 2017; Graham et al., 2013; Graham, Meindl, Beall, Johnson, & Zhang, 2016; Haidt, 2007; Miller, 2008). Evidence from neuroimaging studies indicate that moral judgment generally involves a wide network of brain areas on both medial and lateral brain surfaces (Bzdok et al., 2012). Brain regions consistently shown to be related to moral reasoning include the Orbitofrontal Cortex, Insula, Amygdala, Cingulate Cortex, and Precuneus as well as the Temporoparietal Junction, Angular Gyrus, Pallidum and Temporal Pole (for a review, see: Boccia et al., 2017; Sevinc & Spreng, 2014). Additionally, different moral domains seem to be associated with distinct neural networks (Chakroff et al., 2015; Lewis, Kanai, Bates, & Rees, 2012; Parkinson et al., 2011; White et al., 2017).

This variety is in line with the moral foundations theory, which posits that the human moral system is not unitary but rather based on a set of distinct innate intuitions (foundations), such as disgust towards impurity and contamination, condemnation of causing emotional or physical harm, aversion to dishonesty, disapproval of group disloyalty, and dislike for defying authority and tradition (Graham et al., 2013). In fact, using moral transgression scenarios loosely based on the moral foundations theory as stimuli in an fMRI experiment, Parkinson et al. (2011) demonstrated that different moral transgressions evoked activation in specific brain regions. This research has paved the way for further studies on the neural correlates of distinct moral intuitions.

Given the multiple explanations for the negative perception of food-wasting, it remains unclear how the previously mentioned moral foundations are related to moral judgment of this behavior. This issue was investigated in the current study. We were interested in answering the following questions. (1) Is food-wasting considered immoral in reference to other unequivocally immoral categories, such as moral disgust, harm, and dishonesty? (2) Which of these categories of moral intuitions best capture people’s moral judgment regarding wasting food? We sought to answer these questions using both behavioral and neuroimaging data.

## Methods

To address the research questions using an fMRI experiment, we first needed to design appropriate stimuli. Therefore, in the first phase of our study, we developed a standardized set of affective scenarios while ensuring that each scenario described a behavior belonging to one of the following categories: food-wasting behavior, morally disgusting behavior, emotionally or physically harmful behavior, dishonest behavior, and neutral behavior (see *Pilot Study*). In the main experiment, the participants made moral judgments regarding each scenario while in an fMRI scanner. In the post-imaging procedure, we collected detailed responses on the moral appraisal of each scenario and the affective experience of reading each scenario according to the dimensions of emotional valence, arousal, and imageability, as well as data on the environmental attitudes, empathy and demographics of the participants. The assessments of the food-wasting scenarios were then compared to other categories. The study complied with the Declaration of Helsinki on Biomedical Research Involving Human Subjects and was approved by the institutional review board of the relevant university. All participants provided informed consent prior to participation.

### Pilot Study

We conducted a pilot study to create appropriate stimuli for the main experiment. In the first step, we created 30 scenarios in each of three categories corresponding to moral foundations theory (Graham et al., 2013): disgusting, harmful, dishonest, and 30 food-wasting and 60 neutral scenarios. All scenarios were between 28 and 62 words long (*M* = 43.56, *SD* = 6.64) and started with the name of the protagonist with an even division between female and male names. Approximately 30% of the scenarios were based on scenarios from Parkinson et al. (2012). We translated and culturally adjusted all scenarios that we deemed in line with the descriptions of intuitions of moral disgust, harm, and dishonesty according to the moral foundations theory (Graham et al., 2013) and excluded the scenarios that seemed too unusual (e.g., sex with a dead coyote). The remaining scenarios were created independently by the authors based on the descriptions of moral intuitions as posited by the moral foundations theory (Graham et al., 2013). Scenarios regarding food-wasting behavior presented throwing away good edible food, which was not spoiled or contaminated in any way.

To obtain sets of scenarios that would reliably pertain to the designated categories (3 types of moral foundation, food-wasting, and neutral scenarios), we performed large scale online testing of the created scenarios involving 506 volunteers (367 women, 133 men; 6 participants did not specify their gender; mean age = 26.54, *SD* = 7.92). The pilot study was posted on our own Internet domain with the name of the study in the web address. The study participants were solicited on social media and the university campus. Data collection was anonymous. Through clicking the “Start the procedure” button, the participants acknowledged that they had read and understood the purpose of the study.

Before the start of the assessment procedure, the participants were instructed that they would read a set of 15 scenarios and make several ratings of each one. The participants were informed that the scenarios described real events. The participants were asked to rate the scenarios according to the following questions and instructions:

1. Is the act morally wrong? [*This question does not ask whether the described behavior is or should be legal. It also does not ask whether you are capable of such behavior or whether you would advise such behavior to a friend. The question is only whether you consider the described behavior as immoral. Assume that you only know the given circumstances.*]
2. Is the act harmful? [Does this behavior result in physical or emotional harm to other people?]
3. Is the act dishonest? [Did anyone get cheated, unequally treated or have something stolen as a result of this behavior?]
4. Is the act disgusting? [Does this behavior fill you with disgust or revulsion?]

The participants answered each question on a scale from 0 to 7, where 0 meant “definitely not” and 7 meant “definitely yes”. Additionally, they marked their reactions on nine-point pictorial “valence” and “arousal” Self-Assessment Manikin (SAM) scales (Bradley & Lang, 1994). The assessment ranged from 1 to 9 (“very negative”/ “very positive” in case of valence, and “not at all aroused”/ “very much aroused” in case of arousal). Every participant rated 15 randomly selected scenarios from the set of 180 scenarios. The entire procedure required approximately 15 minutes to complete. Each item was rated approximately 30 times by 30 different participants.

Scenarios were considered acceptable for use in the main study if they met the following criteria. In the first step, every moral transgression scenario had to be considered immoral (i.e., its mean rating on the immorality scale was above 3). In the second step, it had to be perceived as representative of its intended moral domain (i.e., the mean ratings on the three moral transgression category scales were the highest for the intended category). For the food-wasting items, only the first criterion was applied. A neutral scenario was deemed acceptable if it was not considered immoral; i.e., its mean rating on each of moral transgression scales did not exceed .5.

The results of the pilot testing yielded 12 suitable scenarios in each of the moral transgression categories and the food-wasting category, and 40 neutral scenarios. All final scenarios were between 40 and 48 words long (*M* = 44.36, *SD* = 2.01) and started with the name of the protagonist (e.g. “George is on a business trip in India and leaves the hotel for dinner at street vendors. The dishes are so tasty and cheap that he buys a few bags full of food to try several different dishes. After a while, he is already full, so he throws two full bags of food into the trash.”). We modified the scenarios so that the numbers of female and male protagonists were equal. The scenarios selected for the fMRI study did not significantly differ in length between categories (mean scenario lengths for all five categories were approximately 44 words; *F*(4, 83) = .33, *ns*). For the contents of scenarios in different categories, see the supplementary material.

### fMRI study

#### Participants

Fifty naive adults aged 20-35 years (*M* = 26.82, *SD* = 3.94, 25 men) took part in the study. All participants were right-handed residents of a European city (students and/or working people) with normal or corrected-to-normal vision. During the recruitment process, we used the adapted version of Early Life Stress Questionnaire (Mcfarlane et al., 2005) to exclude individuals who might have experienced trauma and could therefore feel overly uncomfortable by the explicit content of certain scenarios used in the study. All participants were instructed to come to the study no more than two hours after a meal to avoid the possibility that hunger confounds moral appraisal of wasting food (Simmons et al., 2013). All subjects received a financial reward.

#### Procedure

Before scanning, the experimenter obtained informed consent and explained the task to the participant. Participants were told that they would read multiple scenarios describing the actions of different people. They were asked to consider whether each act they would read about was morally wrong and, if so, whether it was so because it was harmful, dishonest, or disgusting. They were also given definitions for these categories (see *Pilot Study*). Next, the participants were instructed that some of the presented scenarios would have to be evaluated using response pads. In this case, after viewing the fixation cross which followed a scenario, they would be prompted to indicate on four independent seven-point scales to what extent they consider the described act to be morally wrong, harmful, dishonest, and disgusting, respectively and to mark their emotional reactions on two independent nine-point pictorial “valence” and “arousal” SAM scales (Bradley & Lang, 1994). The scales are described in detail in Section *Pilot Study*.

Prior to the start of the fMRI procedure, each participant underwent a short training session on a computer, during which he or she read detailed instructions regarding the procedure and rated 4 exemplary scenarios (used exclusively in the training session) on the six previously described scales using the same response pads as employed later in the scanner. Participants were taught how to move left and right on the scales and to confirm their answer with a specific button on the response pad. The response procedure was introduced to ensure that participants would remain alert and read the scenarios.

The fMRI procedure was divided into three sessions. In the first session, the participants read 30 scenarios while 29 scenarios were presented in each of the remaining two sessions. These sessions consisted of 4 scenarios from each moral transgression category, 4 associated with food-wasting, and 13 (in the 1st session) or 12 (in the 2nd and 3rd session) neutral scenarios. The order of the types of moral transgression and food-wasting scenarios was pseudorandomized, and they were always separated with neutral scenarios. In every session, 4 scenarios (1 per moral transgression category) were to be rated using the response pads. In total, there were 12 scenarios with rating screens per subject, 3 in food-wasting and each moral transgression category. Between reading the scenarios, participants viewed a fixation cross for periods of either 8, 10, or 12 seconds. A schematic representation of a single trial with response scales is presented in Figure 2. The entire fMRI procedure required approximately 50 minutes.

**Fig 2.2.**
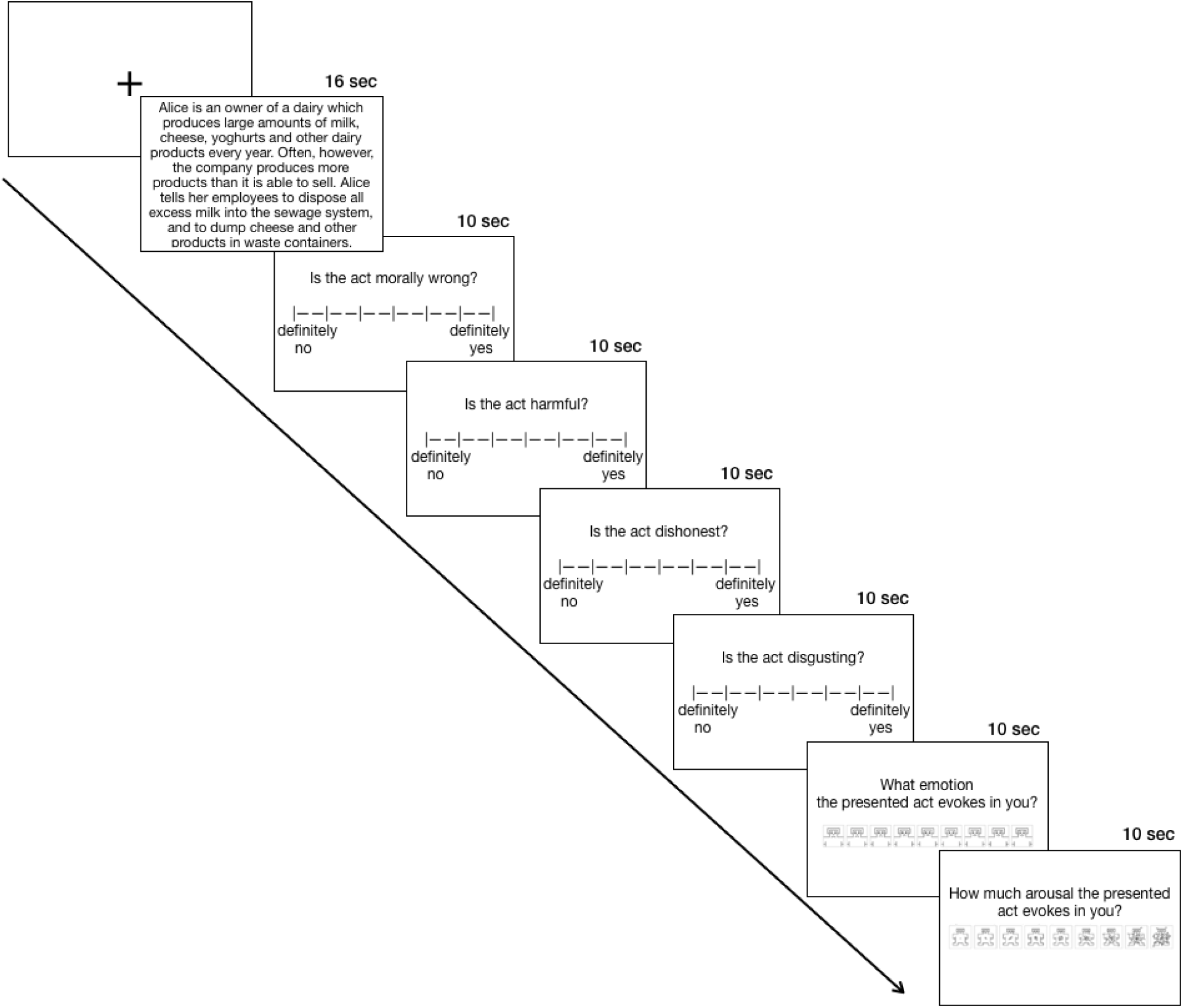
Schematic representation of a single trial with response scales. Participants viewed a fixation cross between trials for periods of either 8, 10, or 12 seconds. Each trial began with a scenario presentation that lasted 16 s. Then, participants were either prompted to only consider in their minds whether the act was morally wrong and if so whether it was so because it was harmful, dishonest, or disgusting or to indicate their answers on four 7-point “moral transgression scales” (1. “Is the act morally wrong?” 2. “Is the act harmful?” 3. “Is the act dishonest?” 4. “Is the act disgusting?”) and two 9-point SAM “emotions scales” (1. “What emotion the presented act evokes in you?” and 2. “How much arousal the presented act evokes in you?”). Each response screen was presented for 10 seconds. In total, there were 12 scenarios with rating screens, 3 for each moral transgression category (disgust, harm, dishonesty) and 3 for food-wasting.

#### fMRI data acquisition

Magnetic resonance imaging data were acquired using a 3T Siemens MAGNETOM Trio system (Siemens Medical Solutions) equipped with a 12-channel head coil. Within a single scanning session, the following images were acquired: structural localizer image, structural T1-weighted image (TR: 2530 ms, TE: 3.32 ms, flip angle: 7°, voxel size: 1 × 1 × 1 mm, field of view: 256 mm), field map magnitude image (TR: 800 ms, TE: 4.5 ms/6.96 ms flip angle: 60°, voxel size: 3 × 3 × 3 mm, field of view: 216 mm), 3 series of functional EPI images (TR: 2500 ms, TE: 28 ms, flip angle: 80°, voxel size: 3 × 3 × 3 mm, field of view: 216 mm, measurements: 413 in the first and 403 in the second and third sessions).

#### fMRI Preprocessing

DICOM series were converted to NIfTI with dcm2niix v. 1.0.20171215 (https://github.com/rordenlab/dcm2niix) using the Horos BIDS Output Extension (https://github.com/mslw/horos-bids-output) as a wrapper. Spatial preprocessing was performed using Statistical Parametric Mapping (SPM12, http://www.fil.ion.ucl.ac.uk/spm/). Functional images were corrected for distortions related to magnetic field inhomogeneity; raw images were corrected for motion by realignment to the first acquired image and signal variance related to movement in magnetic field was corrected by fieldmap unwarping. Prior to normalization, structural images were co-registered to the mean functional image. Functional images were then normalized to the MNI space and re-sliced to 2 × 2 × 2 mm. Finally, images were spatially smoothed with a 6 mm FWHM Gaussian kernel.

#### fMRI data analysis

Normalized, smoothed images were used to compute a 1^st^ level general linear model (GLM) separately for each subject. For each subject, the GLM consisted of three sessions, each containing five regressors of interest (denoting food-wasting, disgusting, harmful, dishonest, and neutral scenarios). The rating periods, during which participants were asked to make moral judgement of the presented scenario, were entered into the GLM as a covariate of no interest. All the regressors related to the course of the experiment were convolved with the standard haemodynamic response function implemented in SPM12 software (https://www.fil.ion.ucl.ac.uk/spm/software/spm12/). Additionally, 6 head-motion parameters and regressors representing each of three scanning sessions were entered into the GLM as covariates of no interest. A standard 128 sec high-pass filter was applied to the data during model estimation. The 2^nd^ level (or group) analyses were computed using a conjunction approach implemented in SPM12 as well as using a full factorial 1×4 model based on 1^st^ level contrasts where food-wasting and moral transgression conditions were contrasted with the neutral condition. Only results that survived FWE correction (*p* < .05) are presented. Peaks of activation were labeled using the Harvard-Oxford atlas (Desikan et al., 2006), unless stated differently in the text.

#### Behavioral study

After leaving the scanner, participants were invited to a separate room where they completed a set of tasks using a computer program designed for the study. In the first part, we assessed the participants’ degree of recall of the scenarios from the fMRI procedure using a 16-item short memory test concerning the contents of selected scenarios. We performed this step to assess the extent to which participants were focused on reading the scenarios in the scanner.

The second part of the post-imaging procedure was identical to the task presented in the scanner where participants had to use response pads to indicate their answers. In the post-imaging procedure, we added an “imageability” scale to assess how easy it was to imagine the act. The scale ranged from 1 - “I could not imagine it at all” to 9 - “I could completely imagine it”. A printout of the definitions of the moral domains used across the study was always available on the computer desk. In this way, the participants assessed all 88 scenarios used in the fMRI procedure.

Taking into consideration that moral decisions are influenced by individual differences, for example, in empathy (Yoder & Decety, 2018), we wished to assess the levels of this trait in our participants as well as their degree of pro-environmental orientation. We assumed that empathy would be linked to harsher moral appraisals of all types of transgression, whereas a stronger environmental orientation would be related to harsher moral judgment regarding wasting food. We collected information on the participants’ level of empathy using the 28-item Empathic Sensitiveness Scale (Kaźmierczak, Plopa, & Retowski, 2007) and their degree of pro-environmental orientation using the adapted version of the revised 15-item New Environmental Paradigm (Dunlap, Van Liere, Mertig, & Jones, 2000). The subjects responded to the statements in the questionnaires using a 5-point Likert scale. In addition to the age and sex of each participant, we also collected information on the participants’ socioeconomic status (SES) using the MacArthur Scale of Subjective Social Status, which is a reliable tool to measure subjective social status using a numbered stepladder image (Adler, Epel, Castellazzo, & Ickovics, 2000). The entire post-imaging procedure required approximately one hour.

## Results

All behavioral data collected in this study are included in the supplementary material. The neuroimaging datasets generated and/or analyzed during the current study are available from the corresponding author upon request.

### Behavioral Data

Behavioral data analysis was performed using the R language for statistical computing version 3.6.0 (R Core Team, 2019). We used the “nlme” package (Pinheiro, Bates, DebRoy, Sarkar & R Core Team, 2017) to perform multilevel modelling on the within-subject data. Posthoc analysis was performed by means of Tukey tests using the “multcomp” package (Hothorn, Bretz, & Westfall, 2008).

#### Morality and moral transgression types

The results of multilevel modelling revealed significant differences in evaluating scenarios as morally wrong across categories (χ2(8) = 475.82, *p* < .001). The results of post-hoc analysis showed that harmful scenarios were rated as significantly more immoral than all other categories of scenario (harmful – food-wasting: *b* = .98, *p* < .001, harmful - disgusting: *b* = .15, *p* < .001, harmful - dishonest: *b* = .44, *p* < .05, harmful - neutral: *b* = 5.41, *p* < .001), and neutral scenarios were rated as significantly less immoral than all other categories of scenario (neutral - food-wasting: *b* = -4.43, *p* < .001, neutral - disgusting: *b* = - 4.37, *p* < .001, neutral - dishonest: *b* = -4.98, *p* < .001). Dishonest scenarios also differed from all other categories (dishonest - food-wasting: *b* = .55, *p* < .01, dishonest - disgusting: *b* = .15, *p* < .001). There were no statistically significant differences between the ratings of immorality of food-wasting and disgusting scenarios. See Figure 3.1. for details.

**Fig 3.1.**
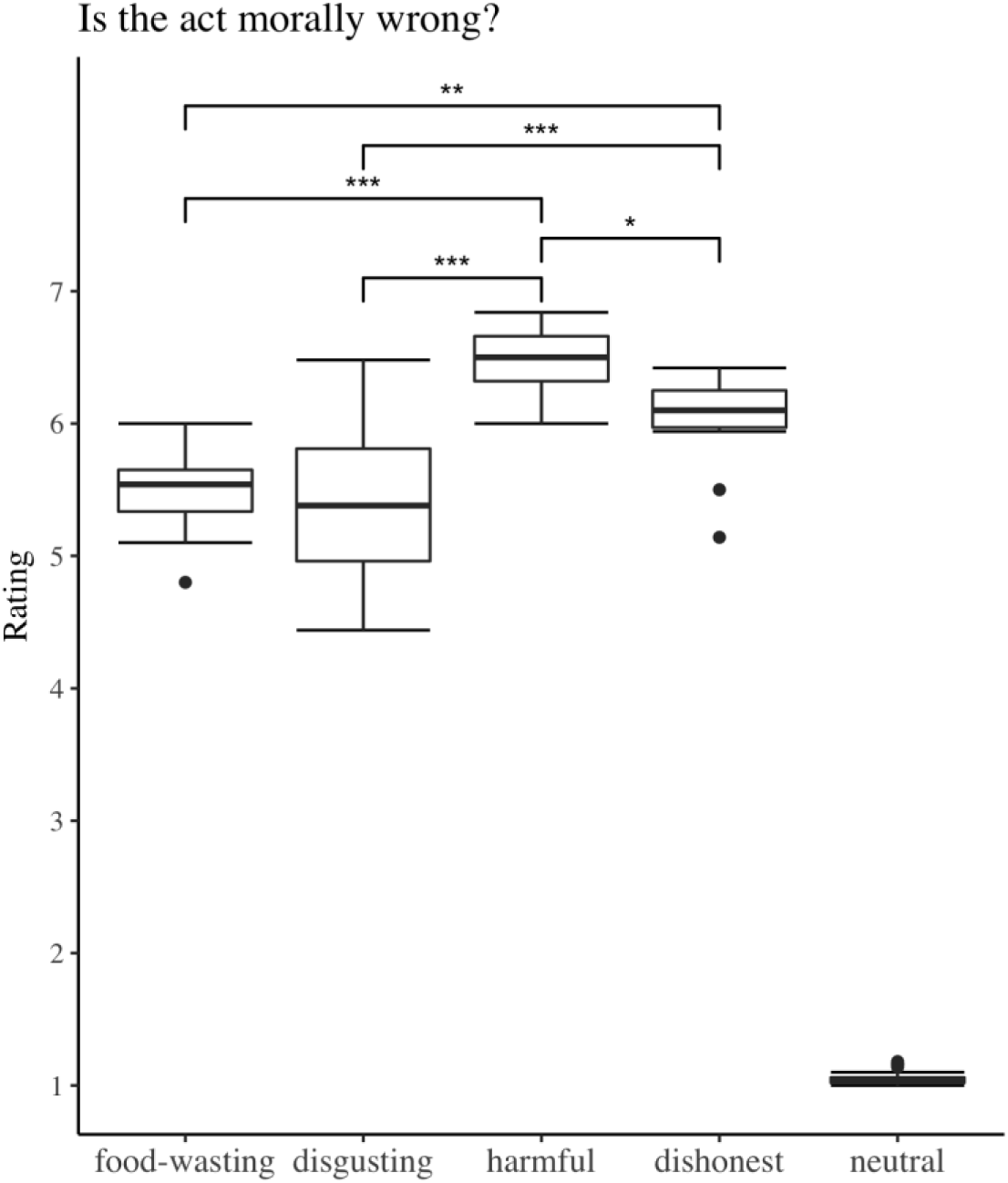
Post-imaging mean ratings of moral wrongness across different categories of scenario. Neutral scenarios were significantly less immoral than all other categories. Significance codes: ‘***’: *p* < .001; ‘**’: *p* < .01. ‘*’: *p* < .05.

Further multilevel modelling indicated that the scenarios were classified into the intended moral transgression categories. There were significant differences in ratings of different categories of scenario across three moral categories (for disgusting scenarios: χ2(2) = 95.98, *p* < .001; for harmful scenarios: χ2(2) = 70.39, *p* < .001; for dishonest scenarios: χ2(2) = 98.76, *p* < .001). Post-hoc tests revealed that disgusting scenarios were rated as significantly more disgusting than harmful (*b* = 1.58, *p* < .001) or dishonest (*b* = 2.13, *p* < .001), and that harmful scenarios were rated as significantly more harmful than disgusting (*b* = 1.08, *p* < .001) or dishonest (*b* = 2.11, *p* < .001). Dishonest items were rated as significantly more dishonest than disgusting (*b* = 2.53, *p* < .001) or harmful (*b* = 1.74, *p* < .001). Neutral scenarios were rated as neither disgusting (*M* = 1.04) nor harmful (*M* = 1.05) nor dishonest (*M* = 1.04). There were no significant differences among ratings of categories across neutral scenarios (χ2(2) = 2.08, *p* = .35).

In addition, there were significant differences in the ratings of food-wasting scenarios across moral transgression scales (χ2(2) = 24.97, *p* < .001). Post-hoc analyses revealed that food- wasting scenarios were rated as significantly more disgusting than harmful (*b* = .54, *p* < .05) and dishonest (*b* = 1.17, *p* < .001). They were also seen as significantly more harmful than dishonest (*b* = 0.63, *p* < .05). Ratings of harm, dishonesty and disgust across different scenario categories are presented in Figure 3.2

**Fig. 3.2.**
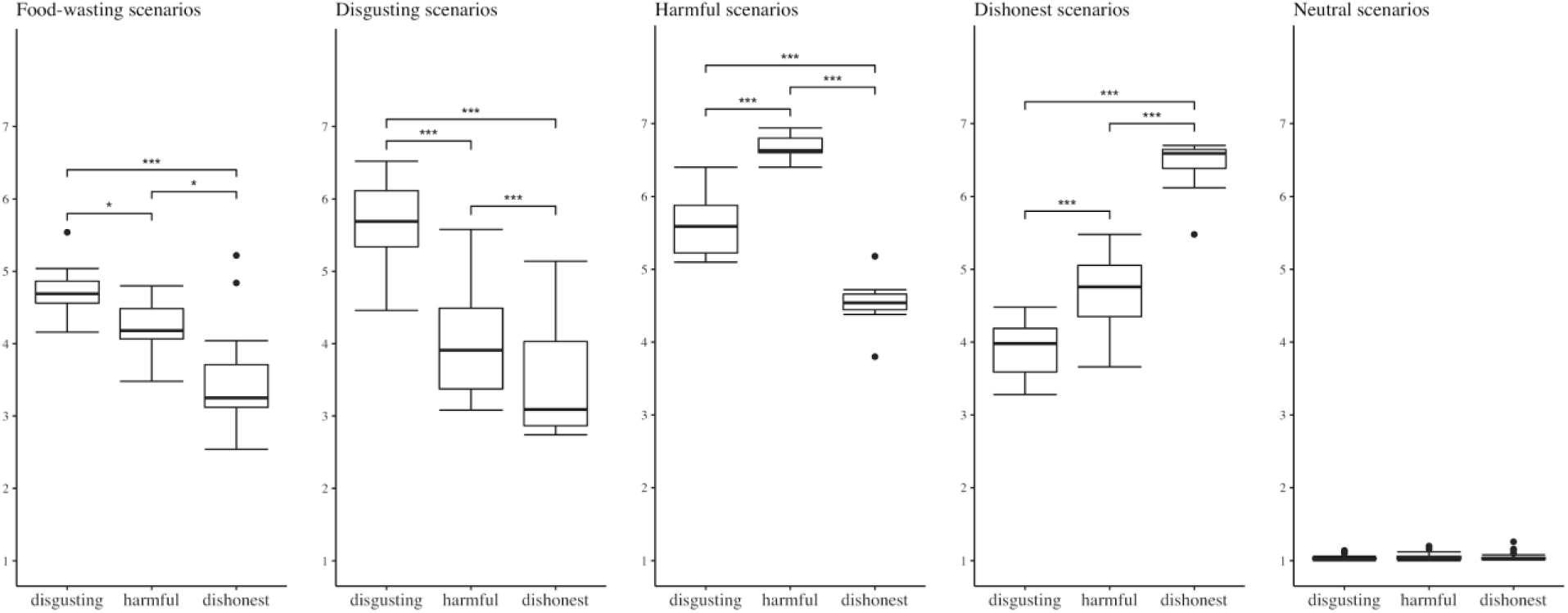
Post-imaging mean ratings of harm, dishonesty and disgust across different categories of scenario. Significance codes: ***: *p* < .001; **: *p* < .01; *: p < .05.

#### Ratings of emotional valence, arousal and imageability

The scenario category had a significant effect on the rating of emotional valence, χ2(8) = 282.43, *p* < .001. Tukey’s post hoc analysis showed that participants rated harmful scenarios as significantly more emotionally negative than all other categories of scenarios (harmful – food-wasting: *b* = -.97, *p* < .001, harmful disgusting: *b* = -1.04, *p* < .001, harmful - dishonest: *b* = -1.26, *p* < .001, harmful - neutral: *b* = - 3.96, *p* < .001). There were no significant differences between other categories except neutral scenarios, which were rated as significantly more positive than all other categories (neutral – food- wasting: *b* = 3.00, *p* < .001, neutral - disgusting; *b* = 2.93, *p* < .001, neutral - dishonest: *b* = 2.70, *p* < .001). See Figure 3.3A for details.

**Fig. 3.3.**
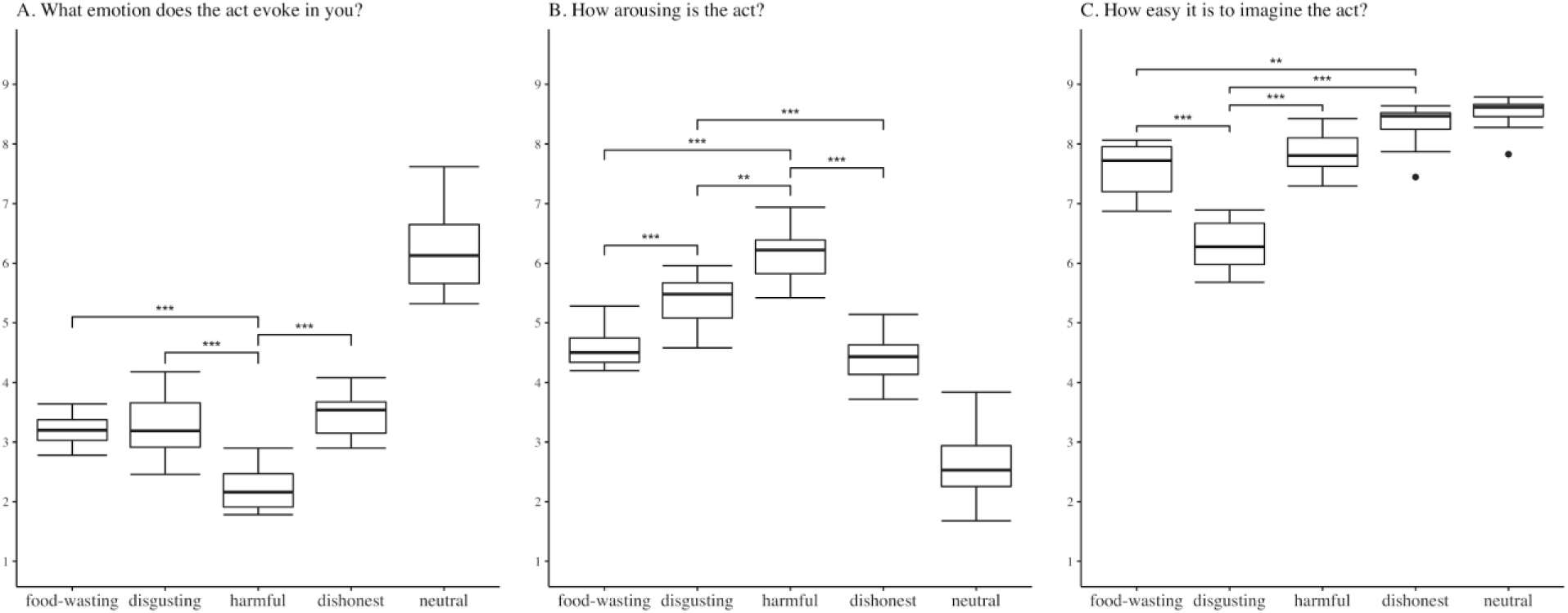
Post-imaging mean ratings of emotional valence, arousal and imageability across different scenario categories. Neutral scenarios were significantly different from other categories in all ratings except for the comparison of imageability ratings with dishonest scenarios. Significance codes: ***: *p* < 0.001; **: *p* < 0.01.

The category of a scenario had a significant impact on the ratings of emotional arousal, χ2(8) = 205.44, *p* < .001. Tukey’s post hoc analysis indicated that participants rated harmful scenarios as significantly more arousing than all other categories of scenario (harmful – food-wasting: *b* = 1.58, *p* < .001, harmful - disgusting: *b* = .76, *p* < .01, harmful - dishonest: *b* = 1.78, *p* < .001, harmful - neutral: *b* = 3.55, *p* < .001). Disgusting scenarios also significantly differed from all other categories (disgusting – food-wasting: *b* = .82, *p* < .001, disgusting - dishonest: *b* = 1.02, *p* < .001, disgusting - neutral: *b* = 2.8, *p* < .001). Additionally, food-wasting scenarios were rated as significantly more arousing than neutral scenarios (*b* = 1.98, *p* < .001), similarly to dishonest scenarios (dishonest - neutral: *b* = 1.78, *p* < .001). For details, see Figure 3.3B.

Category of scenario had a significant impact on the ratings of imageability of presented acts, χ2(8) = 109.13, *p* < .001. Tukey’s post hoc analysis indicated that participants rated disgusting scenarios as significantly more difficult to imagine than all other categories of scenario [disgusting- food-wasting (*b* = -1.28, *p* < .001), disgusting - harmful (*b* = -1.56, *p* < .001), disgusting - dishonest (*b* = -2.01, *p* < .001), disgusting - neutral (*b* = -2.24, *p* < .001)]. Food-wasting scenarios were significantly more difficult to imagine than dishonest (*b* = -.72, *p* < .01) and neutral scenarios (*b* = -.96, *p* < .001). Additionally, harmful scenarios were significantly more difficult to imagine than neutral scenarios (*b* = .69, *p* < .01). Imageability ratings across different scenario categories are presented graphically in Figure 3.3C.

#### Correlational analysis

Both the Empathic Sensitiveness Scale (ESS) and the revised New Environmental Paradigm scale (NEP_R) had good internal consistency as measured by Cronbach’s coefficient alpha (for ESS α = .82; for NEP α = .8). To inspect whether there was a relationship between particular ratings of different types of scenario and ESS and NEP-R as well as demographic variables, such as age and SES, we computed Spearman’s rank correlation coefficients with Bonferroni correction for mean ratings of moral wrongness, disgust, harm, dishonesty, emotional valence, arousal, and imageability across different categories of scenario. In light of our hypotheses, we found no significant relationships between these variables.

#### Participants’ attention

To control for participant attention when judging scenarios in the scanner, we recorded the participant’s levels of recalling these scenarios at the beginning of the post-imaging procedure. All participants scored significantly above chance (*M* = 88.5%, *min* = 72%). There were significant differences in memory performance between moral transgression categories, with disgusting scenarios being recalled significantly worse than all other types of scenarios (see Supplementary Figure 1). Wilcoxon rank sum tests with Bonferroni correction comparing ratings of moral wrongness, disgust, harm, dishonesty, as well as emotional valence and arousal between the fMRI sessions and the post-imaging assessments exhibited no significant differences between these two rating tasks.

### fMRI study results

To identify the neural correlates of the moral judgment of food-wasting behavior, we conducted four sets of analyses. First, we compared brain activation when making moral judgments regarding scenarios concerning wasting food, and each category of moral transgression with brain activation regarding neutral scenarios (section *Moral transgression scenarios compared with neutral scenarios*). Second, to embed the subsequent analyses in the existing literature, we performed a conjuntion analysis of the disgusting, harmful, and dishonest scenarios (each contrasted with neutral scenarios to exclude information regarding semantic processing of the presented stimuli) (section *Common neural network of moral judgment*). Next, to establish how food-wasting scenarios relate to other moral transgression categories, we performed three sets of conjunction analysis pooling together food-wasting scenarios and each moral transgression category separately (again, each contrasted with neutral scenarios). As we were specifically interested in how food- wasting relates to moral disgust, harm, and dishonesty, we applied a mask that excluded the common neural network of these moral categories from the results of these conjunctions (section *Conjunction analysis*). In the last step, to determine what distinguishes moral appraisal of food- wasting behavior from other types of moral intuitions, we performed a differential analysis involving a contrast of food-wasting scenarios with disgusting, harmful, and dishonest moral transgressions pooled together (each contrasted with neutral scenarios) (section *Food-wasting - a differential analysis*).

#### Moral transgression scenarios compared with neutral scenarios

The judgment of scenarios involving wasting food compared with neutral scenarios indicated increased bilateral activity in a wide spreading brain network including the Frontal Gyrus, Orbitofrontal Cortex, Amygdala, Thalamus, and Caudate, as well as increased activity in the Precuneus, Cingulate Gyrus, Angular Gyrus, Lingual Gyrus, Pallidum, and Parahippocampal Gyrus in the left hemisphere, and the right Middle Temporal Gyrus. For details, see Figure 3.4 and supplementary Table 1.

**Fig 3.4.**
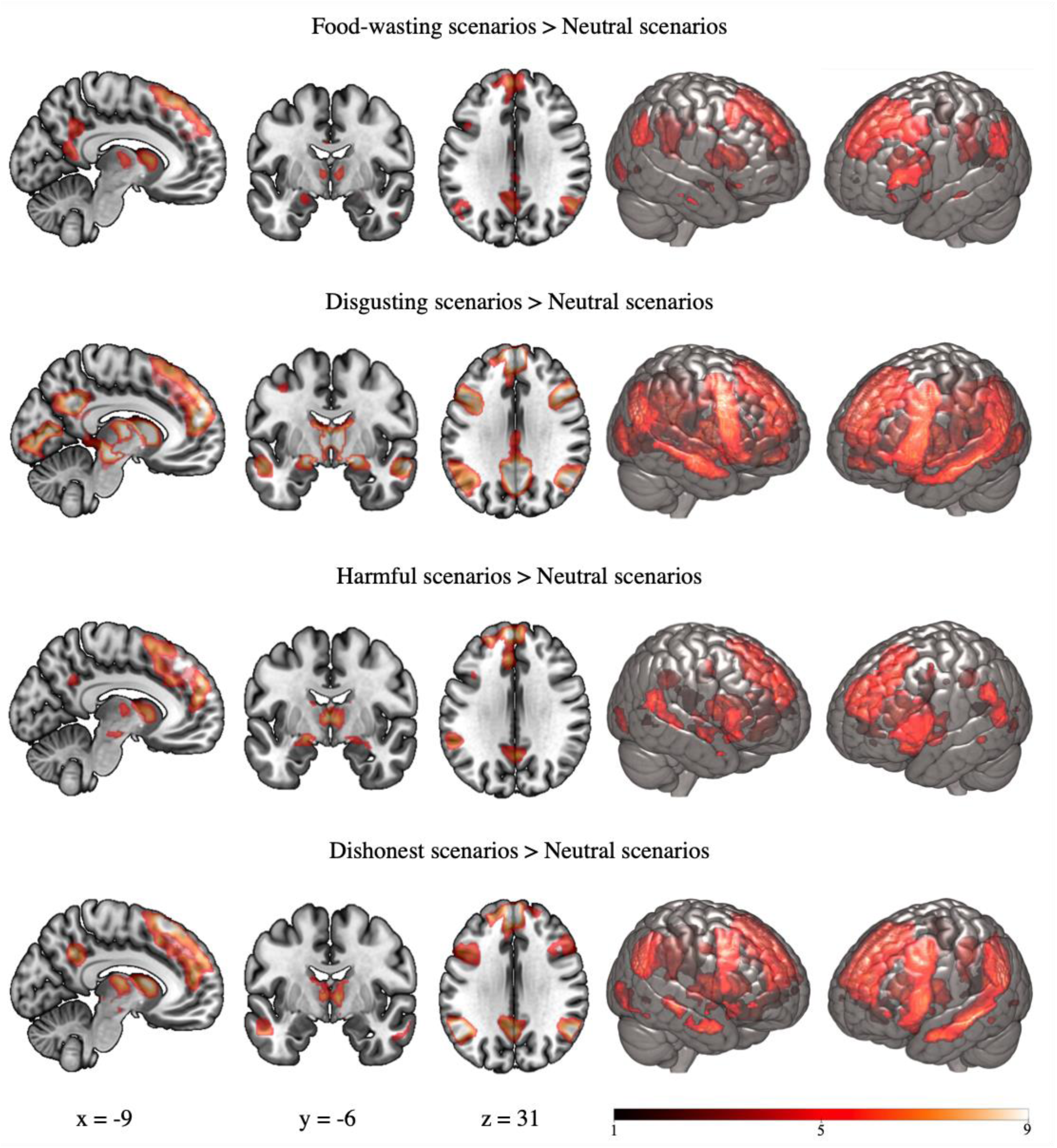
Brain regions revealing significantly greater activation for food-wasting, disgusting, harmful and dishonest moral transgressions than for the neutral scenarios baseline. Results FWE-corrected at *p* < .05.

Moral appraisal of disgusting scenarios compared with neutral scenarios revealed increased activity in the Precuneus Cortex, Frontal Gyrus, Middle Temporal Gyrus, and Temporal Pole in the right hemisphere, as well as the Orbitofrontal Cortex, Cingulate Gyrus and Cerebellum bilaterally. See Figure 3.4 and supplementary Table 1 for details.

The contrast between harmful and neutral scenarios revealed increased activity bilaterally in, among other structures, the Frontal Gyrus, Amygdala, Orbitofrontal Cortex, Supramarginal Gyrus, Middle Temporal Gyrus, Fusiform Gyrus, as well as increased activity in the Insular Cortex and Cingulate Gyrus in the left hemisphere, as well as the right Angular Gyrus. For details, see Figure 3.4 and supplementary Table 1.

Moral judgment of dishonest scenarios compared with neutral scenarios revealed increased activity in the Frontal Gyrus, Angular Gyrus and Middle Temporal Gyrus bilaterally, as well as the Supramarginal Gyrus, Orbitofrontal Cortex, Insular Cortex, and Fusiform Gyrus in the right hemisphere. For detailed information, see supplementary Table 1. This information is also presented graphically in Figure 3.4.

#### Common neural network of moral judgment

The conjunction of disgusting, harmful and dishonest scenarios relative to neutral scenarios revealed increased activity in the brain network including the Frontal Gyrus, Cingulate Gyrus, Supramarginal Gyrus, and Insular Cortex in the left hemisphere, the Precuneus and Middle Temporal Gyrus in the right hemisphere, as well as the Orbitofrontal Cortex, Angular Gyrus, Thalamus and Basal Ganglia bilaterally. For details, see Figure 3.5 and supplementary Table 2.

**Fig 3.5.**
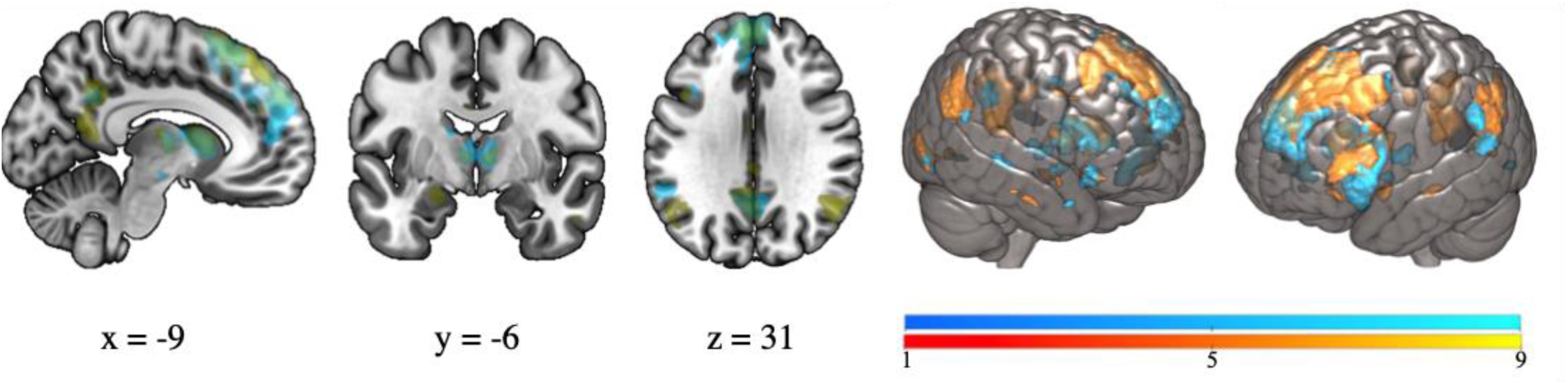
Common activation of moral judgment of disgusting, harmful and dishonest scenarios (in blue) and activation of moral judgment of food-wasting scenarios over neutral scenarios (in yellow). Results FWE-corrected at *p* < .05.

#### Conjunction analysis

The judgment of scenarios involving wasting food and disgusting scenarios indicated increased common activity in a number of structures, including the Orbitofrontal and Cingulate Cortices in the right hemisphere, the left Angular Gyrus, as well as the Amygdalae and dorsal lateral Prefrontal Cortex in both hemispheres. For details, see Figure 3.6 and supplementary Table 3.

**Fig 3.6.**
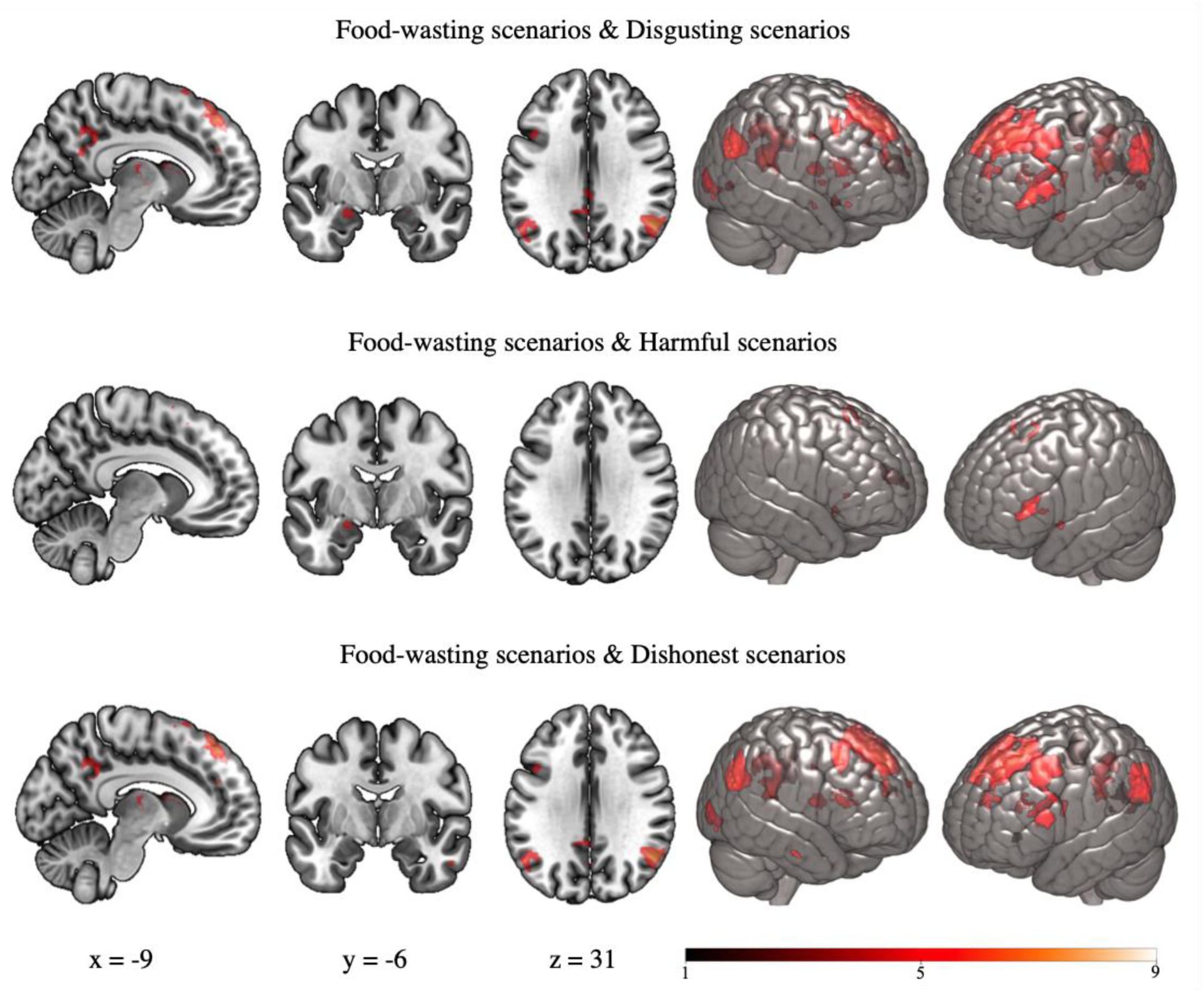
Common activations of moral judgment of food-wasting and disgusting scenarios, food- wasting and harmful scenarios, as well as food-wasting and dishonest scenarios, each masked exclusively with the conjucntion of moral judgment of disgusting, harmful and dishonest scenarios (“common neural network of moral judgment”). Results FWE-corrected at *p* < .05.

Moral appraisal of food-wasting and harmful scenarios revealed increased activity in the bilateral Supplementary Motor Area, as well as Putamen, the Amygdala and dorsomedial Prefrontal Cortex in the left hemisphere. See Figure 3.6 and supplementary Table 3 for details.

The conjunction of food-wasting and dishonest scenarios revealed increased activity in the posterior Cingulate Cortex, Angular Gyrus and Precuneus in the left hemisphere as well as bilateral dorsomedial and ventrolateral Prefrontal Cortex. See Figure 3.6 and supplementary Table 3.

#### Food-wasting - a differential analysis

To establish what distinguishes moral appraisal of scenarios on food-wasting from other types of moral transgression, we contrasted food-wasting scenarios with disgusting, harmful, and dishonest scenarios pooled together. We observed increased activity, among other structures, in the left Insula, bilateral Orbitofrontal Cortex and the right dorsomedial Prefrontal Cortex. For detailed information, see supplementary Table 4. This information is also presented graphically in Figure 3.7.

**Fig 3.7.**
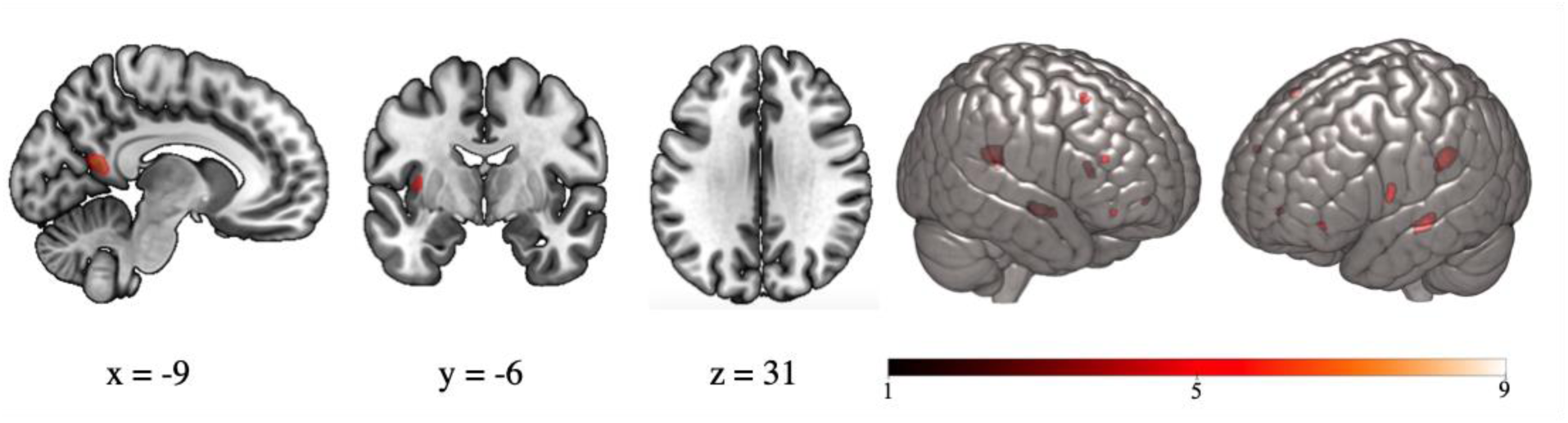
Brain regions showing significant activation for the contrast of food-wasting scenarios against disgusting, harmful, and dishonest scenarios pooled together. Results FWE-corrected at *p* < .05.

## Discussion

The current study sought to examine whether food-wasting is considered immoral in reference to the established moral categories such as moral disgust, harm, and dishonesty. We were also interested in indicating how people’s moral judgment regarding wasting food relates to these three categories of moral intuitions. We are the first to rely on both behavioral and neuroimaging data to study the moral judgment of this harmful phenomenon. Since the existing research indicates that framing persuasive messages referring to moral judgment may be an effective tool to motivate behavior change (Amin et al., 2017), understanding the motives underlying people’s views that throwing away food “is not the right thing to do” may be used in designing successful interventions to tackle this problem.

### Behavioral results

Behavioral analysis showed that food-wasting scenarios were considered immoral to a degree similar to general immorality ratings of disgusting scenarios. Moreover, looking into the ratings in terms of moral categories, food-wasting scenarios were considered primarily morally disgusting and harmful. Hence, on the behavioral level, moral judgment of food-wasting behavior was most related to moral judgment of disgusting behavior.

Food-wasting scenarios were seen as having negative emotional valence, similar in degree to morally disgusting and dishonest scenarios, but not as negative as harmful scenarios. At the same time, scenarios regarding wasting food were moderately arousing. This affective information regarding food-wasting behavior is important in the light of the role of emotion in pro-environmental communication (Chapman, Lickel, & Markowitz, 2017). It has been demonstrated, for example, that experiencing negative affect is related to pro-environmental behaviors (Leviston & Walker, 2012) and support for pro-environmental policies (Smith & Leiserowitz, 2014).

Additionally, moral transgressions seem to always be the source of negative emotions (Avramova & Inbar, 2013). Hence, the perceived negative emotions in the wake of reading food- wasting scenarios confirm that, in people’s view, it is indeed wrong to throw away food. Our results are consistent with other studies suggesting that food-wasting is perceived as a moral problem (Graham-Rowe et al., 2015; Misiak et al., 2018) and expands this conclusion to an industrialized and developed population. As mentioned before, Misiak and colleagues (2018) hypothesized that moral judgments about food-wasting may serve as an adaptation to harsh ecologies. Our findings suggest that moral concerns about food-wasting are observed regardless of the environmental demands.

Scenarios across different categories in our study were seen as relatively easy to imagine. It is important to control this variable, because the degree of imageability of abstract stimuli, such as words, affects behavioral and neuroimaging measures (Westbury et al., 2013). At the same time, when considering differences in imageability of different categories of scenarios, food-wasting scenarios were less imageable than all other categories except morally disgusting scenarios. It has been suggested that people might be aversive to imagining disgusting words because of their strong negative connotations (Thibodeau, 2016). Indeed, words eliciting disgust showed consistent processing disadvantage (Briesemeister, Kuchinke, & Jacobs, 2011; Ferré, Haro, & Hinojosa, 2018). Taking into account that food-wasting behavior was considered mostly morally disgusting, its ratings of imageability might be lowered because of its moral association with disgusting behavior.

### Neuroimaging results

Neuroimaging data analysis demonstrated that moral appraisal of food-wasting scenarios relative to neutral scenarios was associated with an increased activity in regions recognized across studies to be involved in moral judgment such as the bilateral dorsal medial Prefrontal Cortex, right Orbitofrontal Cortex, bilateral Amygdala, as well as anterior Cingulate, Lingual, and Inferior Frontal Gyri in the left hemisphere (Boccia et al., 2017; Parkinson et al., 2011; Sevinc & Spreng, 2014). A large part of this widespread network relevant to moral judgment is also engaged in emotional processing (Kragel & LaBar, 2016), along with subcortical structures such as the Thalamus and Basal Ganglia (Pessoa, 2017) which also revealed increased activity when making moral judgment of food-wasting behavior relative to neutral scenarios in our study. At the same time, an increased activity in the dorsolateral Prefrontal Cortex suggests that moral judgment of wasting food was subject to at least some degree of cognitive control that was suggested to mitigate emotional responses to moral transgressions (Greene, Nystrom, Engell, Darley, & Cohen, 2004). Our study is the first to demonstrate on a neurophysiological level that moral judgment of food- wasting behavior is a complex phenomenon that strongly engages people emotionally and to some degree, also cognitively.

Brain activation related to judgment of disgusting, harmful and dishonest scenarios resembled that from previous studies on neural correlates of moral judgment in these distinct domains (Abe et al., 2014; Parkinson et al., 2011; Schaich Borg, Lieberman, & Kiehl, 2008). Hence, the common neural network of moral judgment of disgusting, harmful, and dishonest scenarios revealed in our study overlaps with structures indicated in previous reports on activation common to moral appraisal of various transgression types including the bilateral dorsal and ventral medial Prefrontal Cortex, bilateral Orbitofrontal cortex, Insular Cortex and Supramarginal Gyrus in the left hemisphere, as well as the Middle Temporal Gyrus and Angular Gyrus in the right hemisphere (Boccia et al., 2017; Sevinc & Spreng, 2014).

Unique activations for the conjunction of the moral judgment of food-wasting and disgusting scenarios elicited overlapping increased activation in brain regions conventionally associated with the processing of physical disgust such as the Orbitofrontal Cortex, Striatum, Amygdala, visual areas and structures involved in the default mode network such as the Angular Gyrus, Cingulate Cortex, Precuneus, and medial Frontal Cortex (Pujol et al., 2018). Some of these structures, especially the Orbitofrontal Cortex, Amygdala, Striatum and mediodorsal Thalamus, are generally related to emotional and reward-related processing (Lindquist, Wager, Kober, Bliss-Moreau, & Barrett, 2012; Sescousse, Caldú, Segura, & Dreher, 2013). Interestingly, the overlapping regions in the posterior Orbitofrontal Cortex, medial Striatum and Amygdalae were recognized to be involved in the processing of primary rewards, i.e. rewards which have an innate value and are essential for survival, among which food- and sex-related stimuli are considered classical examples (Kringelbach, 2005; Sescousse et al., 2013). The peak activations in the overlapping regions of the Orbitofrontal Cortex and Paracingulate Gyrus were recognized to be a part of the orbital network that is believed to regulate human consummatory behaviors (Öngür & Price, 2000). The same regions were active when judging morally disgusting scenarios in previous studies (Parkinson et al., 2011; Schaich Borg et al., 2008) suggesting that moral appraisal of both food-wasting and disgusting scenarios relies on the same reward-related processes in the brain. Through representing the affective value of reinforcers, the co-activated orbital network is engaged in decision making and expectation, crucial for regulating the approach and avoidance behaviors that for most of human history were primarily engaged in decisions about food intake (Kringelbach, 2005).

In fact, the emotion of disgust is commonly associated with food-related stimuli, be it sensory factors such as bad smelling food or the sight of moldy leftovers, or the anticipated harmful consequences related to conditioned reactions to noxious food e.g. poisoning (Paul Rozin & Fallon, 1987). Disgust elicitors are most often directly connected to pathogens that have constituted the main selective pressure through human evolution (Fumagalli et al., 2011). However, disgust has a much more profound impact on our lives than mere food-related experiences (H. A. Chapman & Anderson, 2012; D. Jones, 2007; Vicario, Rafal, Martino, & Avenanti, 2017). For example, a large body of research links the emotion of disgust with judgments of moral wrongness (e.g. Eskine, Kacinik, & Prinz, 2011; Tracy, Steckler, & Heltzel, 2019; Tybur, Lieberman, Kurzban, & DeScioli, 2013), especially in the sanctity domain (Wagemans, Brandt, & Zeelenberg, 2018), which according to the Moral Foundations Theory captures moral intuitions about transgressing physical and spiritual purity (Graham et al., 2013). Moreover, sensory and moral disgust seem to share common characteristics on both neural and physiological levels (H. Chapman, Kim, Susskind, & Anderson, 2009; Vicario et al., 2017).

The common ground for sensory and moral disgust is likely to be related to objects or behaviors that decrease one’s fitness (Tybur et al., 2013). On top of causing emotional disgust, behaviors such as coming in contact with biological contaminants or mating with partners of low sexual value have been moralized across cultures (Koleva, Graham, Iyer, Ditto, & Haidt, 2012). Since food is critical for survival, humans developed elaborate food-related behaviors to minimize risks and enhance chances for survival (Rozin, 2001). Following this line of thought, many food-related behaviors are also subject to moral judgment (Lieberman, Tybur, & Latner, 2012; Scott, Inbar, & Rozin, 2016), which is mirrored by the prevalence of food-related taboos across cultures (Meyer-Rochow, 2009) and confirmed both by our behavioral and neuroimaging research. Taking into account that food has not been in surplus for much of human history, the fact that throwing away edible food causes uneasiness is likely to be related to viewing this behavior as reducing fitness, which in turn leads to intuitively judging it as morally disgusting.

Moral judgment of food-wasting and harmful scenarios also revealed an increased activation in several unique areas (absent in morally disgusting and dishonest conditions) including the bilateral Supplementary Motor Area, as well as the Inferior Frontal Gyrus, Putamen and the Amygdala in the left hemisphere. These regions correspond with the neural network involved in empathy (Del Casale et al., 2017; Goerlich-Dobre, Lamm, Pripfl, Habel, & Votinov, 2015; Wu et al., 2018). However, we did not observe activation of the so-called pain matrix (anterior Insula, anterior Cingulate Cortex), which in previous studies consistently showed increased activation while experiencing pain oneself, as well as while watching others experience pain (Lamm, Decety, & Singer, 2011). Nevertheless, the observation of other people being harmed still activated the motor system, which is believed to be an early automatic mechanism of evoking an empathic response (Fabi & Leuthold, 2017; Lamm & Majdandžić, 2015). The shared activation of areas of the empathy network when making moral judgments about food-wasting and harmful scenarios might therefore be associated with perceiving food-wasting as noxious (implicitly for other people or the planet), something similar to seeing people being physically or emotionally harmed.

Alternatively, considering that the comprehension of a word seems to be associated with activation of its articulatory motor program (Pulvermüller, 2005), the content of food-wasting and harmful scenarios might include words processed by overlapping regions in the motor area. It has been shown that action verbs and food nouns elicited similar brain response patterns in the left Inferior Frontal Gyrus and Motor Cortex (Carota, Kriegeskorte, Nili, & Pulvermüller, 2017) which overlap with our results. On the other hand, processing of food-wasting and harmful scenarios led to an increased activation in the left Amygdala and left dorsolateral Prefrontal Cortex, which are functionally connected during emotion regulation (Herwig et al., 2019), suggesting considerable emotional engagement during moral judgment of these scenarios. Since emotion regulation is considered an important component of human empathy (Schipper & Petermann, 2013), the involvement of this limbic-frontal circuitry regulating emotions points to the conclusion that the shared pattern of brain activity during moral judgment of food-wasting and harmful scenarios is rather related to automatic empathic response than language processing.

Increased common activations specific to food-wasting and dishonest scenarios included a network involved in emotional perspective taking, recalling emotional experiences to guide behavior, and integrating emotion into social cognition comprising the medial Prefrontal Cortex, Posterior Cingulate, and Angular gyrus (Britton et al., 2006; Raine & Yang, 2006). Its link to fairness has been demonstrated in a study of psychopathic individuals, in which it was found that these exact structures exhibited reduced activity when judging moral dilemmas in subjects scoring high on the scales of pathologic dishonesty (Glenn, Raine, & Schug, 2009). Moreover, these regions were identified as specifically linked to moral judgment of injustice by Parkinson and colleagues (2011), who attributed their findings to dishonest transgressions requiring more mentalizing than other types of transgressions. However, we observed their activation also for the conjunction of food-wasting and disgusting scenarios, which implies that their part in moral reasoning is broader than only dishonesty appraisal.

The posterior Cingulate Cortex, medial Prefrontal Cortex, Angular Gyrus and Precuneus are the key parts of the default mode network (Utevsky, Smith, & Huettel, 2014) steering internally guided decision-making. Such internally directed processing occurs when individuals refer to their own preferences rather than to external cues in behavior regulation (Leech, Kamourieh, Beckmann, & Sharp, 2011; Nakao, Ohira, & Northoff, 2012), which seems to play a vital role in moral judgment and Theory of Mind (Reniers et al., 2012). These structures were active for the conjunction of food-wasting and disgusting as well as food-wasting and dishonest scenarios suggesting that moral judgment of these behaviors relies on internally guided decision-making to a greater extent than moral appraisal of harmful behavior. Possibly, moral judgment of scenarios presented in these categories was more ambiguous and thus required more internally directed thought than the judgment of emotionally and physically harmful scenarios, which were rated as the most immoral, most negative in valence, and most arousing in the behavioral part of the study. Thus, when judging food-wasting, disgusting, and dishonest behaviors, participants had to deliberately refer to their own value system, perhaps due to the lack of definite ready-to-use responses implied by social norms.

In contrast to disgusting, harmful, and dishonest transgressions pooled together, moral judgment of food-wasting scenarios showed increased activity in regions recognized in the processing of food-related stimuli such as the Insula, Orbitofrontal Cortex and Parahippocampal Gyrus (Chao et al., 2017; Porubská, Veit, Preissl, Fritsche, & Birbaumer, 2006; St-Onge, Sy, Heymsfield, & Hirsch, 2005). Looking into functional specialization of these structures in more detail confirmed this observation. The peak activation of the mid Insula was associated with food both using the Neurosynth automated associations (Yarkoni, Poldrack, Nichols, Van Essen, & Wager, 2011) as well as in our literature review (Avery et al., 2015; Kurth, Zilles, Fox, Laird, & Eickhoff, 2010; Veldhuizen et al., 2011), and this association exceeded primary sensory processing and included visual food cues (van der Laan, De Ridder, Viergever, & Smeets, 2011). Likewise, peak activations of the Orbitofrontal Cortex in this condition were automatically associated with key words “food”, “eating”, and “arousal” in the left hemisphere, and “food”, “reward”, and “emotion regulation” in the right hemisphere (Yarkoni et al., 2011). We suggest that, on top of the areas believed to be underpinning moral reasoning, moral appraisal of food-wasting behavior also engages structures involved in the processing of food-related stimuli. As shown above, these regions are partly interwoven with the neural affective reward system that contributes to moral appraisal of disgusting behavior.

### Limitations and future directions

Our study had several limitations. First, our sample was very homogenous in terms of age, socio-economic status, levels of empathy, and pro-environmental attitudes. For this reason, the correlational analysis did not reveal any links between individual differences and moral and emotional appraisal of moral transgressions. Future studies could differentiate the sample more to track individual differences and their relationships with the moral judgment of wasting food. One interesting variable to take into account would be disgust sensitivity, which was shown to be highly related to moral judgment (Jones & Fitness, 2008; Wagemans et al., 2018). Another limitation to consider is the standardization of stimuli. Similar to other studies on morality based on ad-hoc constructed, text-based stimuli (Boccia et al., 2017), we controlled the length of scenarios, their general construction, and protagonists’ sex. Recently, a report of text stimuli controlled on more dimensions, such as e.g. syntactic structure and complexity, has been published (Clifford, Iyengar, Cabeza, & Sinnott-Armstrong, 2015) allowing for more controlled comparisons and facilitating further studies. Future research could also investigate moral judgment of food-wasting behavior in relation to other moral transgressions relying on a recently published standardized set of images (Crone, Bode, Murawski, & Laham, 2018) or films (McCurrie, Crone, Bigelow, & Laham, 2018), as well as look into neural correlates of moral concerns about waste and inefficiency in general as suggested by Shulman and Mastronarde in Graham and colleague’s work on Moral Foundations Theory (2013).

Since previous studies have demonstrated that emotional valence and arousal predict moral judgment (Szekely & Miu, 2015a; Zhang, Kong, & Li, 2017), one could argue that moral scenarios should also be controlled according to their affective information. However, emotions elicited by transgressions across different moral domains vary in valence and arousal (Szekely & Miu, 2015b). It is important, therefore, to study the range of emotions accompanying moral judgment in different moral domains, rather than to artificially reduce their impact on moral appraisal.

Moral intuitions were recognized to play a key role in motivating morally relevant behavior (Bazerman & Tenbrunsel, 2012; Haidt, 2001). Activating people’s moral intuitions can thus push people into environmentally-relevant action (Markowitz & Shariff, 2012). Moral intuition’s framework has already been used in the study of energy decisions (Sovacool, Heffron, McCauley, & Goldthau, 2016), change of political attitude (Day, Fiske, Downing, & Trail, 2014), and to explore factors underlying vaccine hesitancy (Amin et al., 2017), as well as to predict willingness to make pro-environmental lifestyle changes (Dickinson, McLeod, Bloomfield, & Allred, 2016). Our findings on moral intuitions and emotions which are involved in the appraisal of food-wasting behavior can therefore be applied in persuasive messaging and other behavioral interventions aiming to reduce the problem of food-wasting.

## Conclusion

To our knowledge, we are the first to demonstrate that food-wasting is considered immoral on both behavioral and neuronal level, and that moral judgment regarding wasting food is primarily related to moral judgment of disgusting behavior. The pattern of overlapping brain activations associated with moral judgment of food-wasting and disgusting scenarios shared certain similarities with more general processing of disgusting stimuli. These overlapping brain regions exceeded the common neural network of moral judgment in the dorsomedial and ventrolateral Prefrontal cortex, and comprised areas associated with affective reward-related processing engaged in steering approach/avoidance behaviors and food-intake, as well as the Default Mode Network regulating internally guided decision-making. Moral judgment of wasting food also shared common characteristics with moral appraisal of harmful behavior, suggesting some degree of empathic involvement in judgment of wasting food. At the same time, judgment of food-wasting was different from judgement of other moral transgressions on the neural level with regard to structures related to processing of food-related stimuli. We have also contributed to the growing field of moral (neuro)psychology with our behavioral and neuroimaging data on moral disgust, harm, and dishonesty, which are three categories of moral intuitions already established in the morality literature. Our findings in this regard fall in line with previous research, amplifying our knowledge of the distinct and shared neural correlates of different categories of moral judgment, as well as their emotional and imageability qualities. Lastly, our findings are potentially valuable in the light of moral intuitions’ influence on behavior and may be used in addressing the pressing problem of food-wasting.

## Acknowledgements

The authors were financed by the Polish National Science Centre OPUS grant (2015/19/B/HS6/00331) to Agnieszka Sorokowska. The project was realized with the aid of CePT research infrastructure purchased with funds from the European Regional Development Fund as part of the Innovative Economy Operational Programme, 2007–2013. We would like to thank Filip Gromuł for his invaluable help with the development of the computer program for data collection in the behavioral part of the study and Michał Stefańczyk for assistance during data collection. Many thanks to Paweł Orłowski, Bartosz Kossowski, and Małgorzata Wierzba for their technical support with the development and conduction of this study, as well as to Thomas Custer for proofreading.

