## Supplementary Material for "Wasting food is disgusting: Evidence from behavioral and neuroimaging study of moral judgment of food-wasting behavior"

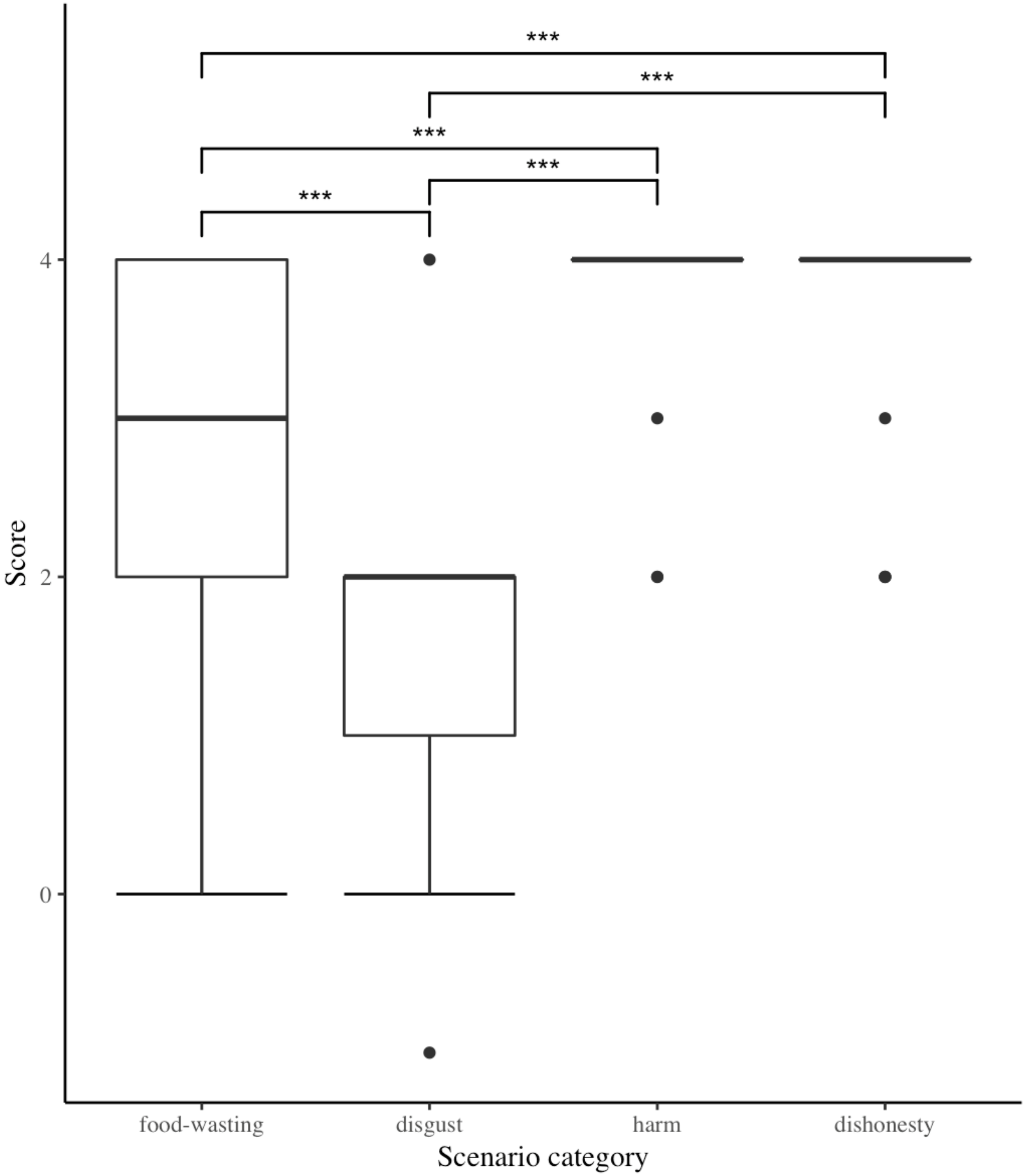

**Supplementary Fig 1.** Differences in memory performance between categories of scenarios.

**Supplementary Table 1**. MNI coordinates for the peak activations (up to 10 peaks per cluster that are at least 8 mm apart, thresholded at 5 voxels) for each moral transgression category versus neutral scenarios (*T* (1, 245) = 4.77, Voxel-wise FWE corrected *p* < .05). Region labels are according to the Harvard-Oxford Atlas.

| **Region Label** | **voxels** | **t-value** | **x** | **y** | **z** |
|---|---|---|---|---|---|
| *Food-wasting scenarios > Neutral scenarios* | | | | | |
| Superior Frontal Gyrus | 4989 | 8.94 | -14 | 32 | 56 |
| | | 8.22 | -16 | 22 | 60 |
| | | 8.09 | 14 | 36 | 52 |
| Frontal Pole | | 8.01 | 12 | 50 | 42 |
| Superior Frontal Gyrus | | 7.82 | 6 | 44 | 50 |
| Frontal Pole | | 7.79 | -8 | 40 | 52 |
| Superior Frontal Gyrus | | 7.78 | -6 | 54 | 34 |
| Middle Frontal Gyrus | | 7.51 | 38 | 22 | 52 |
| | | 7.47 | -30 | 22 | 50 |
| Superior Frontal Gyrus | | 7.30 | -22 | 24 | 50 |
| Right Caudate | 314 | 7.99 | 12 | 10 | 10 |
| | | 6.67 | 14 | 2 | 18 |
| Right Thalamus | | 6.19 | 8 | -4 | 4 |
| | | 5.50 | 10 | -10 | 10 |
| Left Caudate | 386 | 7.96 | -10 | 10 | 8 |
| | | 7.42 | -12 | 4 | 14 |

| | | | | | |
|---|---|---|---|---|---|
| Left Thalamus | | 6.32 | -6 | -8 | 4 |
| Frontal Pole | 560 | 7.51 | -46 | 38 | 10 |
| Inferior Frontal Gyrus, pars triangularis | | 5.77 | -50 | 24 | 8 |
| Lateral Occipital Cortex, superior division | 473 | 7.37 | 54 | -60 | 36 |
| | | 7.18 | 56 | -62 | 28 |
| | 473 | 5.64 | 50 | -58 | 48 |
| Precuneus Cortex | 949 | 7.19 | -2 | -54 | 14 |
| | | 6.93 | 0 | -60 | 34 |
| Cingulate Gyrus, posterior division | | 5.94 | -8 | -52 | 32 |
| Occipital Pole | 206 | 6.69 | 26 | -94 | -4 |
| | | 5.74 | 32 | -90 | -8 |
| Lateral Occipital Cortex, inferior division | | 5.30 | 36 | -88 | 0 |
| Occipital Pole | | 4.98 | 36 | -90 | 8 |
| Lateral Occipital Cortex, superior division | 543 | 6.22 | -42 | -70 | 40 |
| | | 6.22 | -56 | -62 | 24 |
| Angular Gyrus | | 5.56 | -44 | -58 | 34 |
| | | 5.53 | -50 | -58 | 44 |
| Lateral Occipital Cortex, superior division | | 5.26 | -50 | -70 | 32 |
| | | 5.24 | -34 | -74 | 48 |
| Frontal Orbital Cortex | 29 | 6.15 | -28 | 34 | -12 |
| Middle Temporal Gyrus, posterior division | 22 | 5.89 | 54 | -10 | -28 |

| | | | | | |
|---|---|---|---|---|---|
| Left Pallidum | 10 | 5.64 | -16 | 6 | -2 |
| Left Amygdala | 42 | 5.55 | -24 | -6 | -16 |
| Lingual Gyrus | 21 | 5.44 | -4 | -86 | -6 |
| Cingulate Gyrus, posterior division | 46 | 5.27 | -2 | -32 | 34 |
| Cingulate Gyrus, anterior division | 31 | 5.20 | 0 | -14 | 34 |
| | | 5.17 | -4 | -4 | 30 |
| Parahippocampal Gyrus, posterior division | 32 | 5.06 | -30 | -30 | -18 |
| Right Amygdala | 7 | 5.04 | 22 | -4 | -18 |
| Frontal Orbital Cortex | 8 | 5.00 | 44 | 30 | -12 |
| Middle Temporal Gyrus, posterior division | 5 | 4.97 | 60 | -18 | -22 |
| *Disgusting scenarios > Neutral scenarios* | | | | | |
| Precuneus Cortex | 28595 | 17.75 | 2 | -60 | 30 |
| Right Caudate | | 13.44 | 10 | 12 | 6 |
| | | 12.81 | 12 | 6 | 12 |
| Superior Frontal Gyrus | | 12.68 | 0 | 46 | 38 |
| Right Thalamus | | 12.51 | 6 | -4 | 2 |
| Superior Frontal Gyrus | | 12.37 | 2 | 52 | 32 |
| Left Thalamus | | 12.37 | -4 | -6 | 4 |
| Superior Frontal Gyrus | | 12.37 | 2 | 56 | 24 |
| Frontal Orbital Cortex | | 11.56 | -28 | 16 | -18 |
| Left Caudate | | 11.54 | -12 | 4 | 14 |

| | | | | | |
|---|---|---|---|---|---|
| Lateral Occipital Cortex, superior division | 1582 | 12.36 | 54 | -64 | 26 |
| | | 8.96 | 46 | -68 | 42 |
| Middle Temporal Gyrus, temporooccipital part | | 6.43 | 64 | -52 | 12 |
| Middle Frontal Gyrus | 3384 | 9.83 | 56 | 22 | 32 |
| | | 9.61 | 46 | 20 | 30 |
| Frontal Pole | | 9.59 | 38 | 34 | -14 |
| Frontal Orbital Cortex | | 9.40 | 30 | 20 | -16 |
| Middle Frontal Gyrus | | 9.06 | 44 | 18 | 38 |
| Inferior Frontal Gyrus, pars triangularis | | 8.82 | 56 | 26 | 10 |
| Middle Frontal Gyrus | | 8.76 | 44 | 12 | 54 |
| Inferior Frontal Gyrus, pars triangularis | | 8.56 | 48 | 26 | 14 |
| Frontal Orbital Cortex | | 7.29 | 32 | 26 | -4 |
| Inferior Frontal Gyrus, pars triangularis | | 6.72 | 50 | 30 | -4 |
| Middle Temporal Gyrus, anterior division | 1286 | 9.81 | 62 | -4 | -16 |
| Middle Temporal Gyrus, posterior division | | 8.41 | 52 | -10 | -14 |
| Temporal Pole | | 7.85 | 40 | 16 | -30 |
| Middle Temporal Gyrus, posterior division | | 6.44 | 66 | -20 | -8 |
| | | 6.25 | 56 | -26 | -8 |
| | | 6.20 | 60 | -32 | -4 |
| Middle Temporal Gyrus, temporooccipital part | | 5.89 | 66 | -38 | -2 |
| Frontal Medial Cortex | 318 | 8.85 | 0 | 44 | -20 |
| | | 8.34 | 0 | 54 | -16 |

| | | | | | |
|---|---|---|---|---|---|
| Frontal Pole | | 4.95 | 0 | 60 | -2 |
| Cerebellar Lobule* | 61 | 7.03 | 0 | -50 | -20 |
| Cingulate Gyrus, anterior division | 59 | 6.02 | 4 | 34 | 6 |
| | | 4.87 | 0 | 36 | -4 |
| Lateral Occipital Cortex, superior division | 22 | 5.16 | -28 | -74 | 22 |
| Cingulate Gyrus, anterior division | 9 | 5.10 | -2 | 22 | 18 |
| *Harmful scenarios > Neutral scenarios* | | | | | |
| Superior Frontal Gyrus | 3551 | 10.23 | -6 | 54 | 28 |
| Paracingulate Gyrus | | 9.10 | -6 | 20 | 46 |
| | | 8.86 | -6 | 52 | 18 |
| | | 8.09 | -6 | 28 | 34 |
| Superior Frontal Gyrus | | 8.01 | -6 | 20 | 56 |
| Frontal Pole | | 7.83 | -22 | 50 | 30 |
| Superior Frontal Gyrus | | 7.64 | 10 | 16 | 64 |
| | | 7.63 | -6 | 14 | 62 |
| | | 7.19 | -4 | 46 | 42 |
| | | 6.62 | -4 | 36 | 40 |
| Right Caudate | 1606 | 9.09 | 12 | 12 | 6 |
| Left Caudate | | 8.96 | -12 | 12 | 2 |
| Left Putamen | | 8.77 | -16 | 6 | -4 |
| Right Caudate* | | 8.68 | 10 | 4 | 4 |

| | | | | | |
|---|---|---|---|---|---|
| Left Thalamus | 1606 | 8.43 | -2 | -6 | 6 |
| Right Caudate | | 8.22 | 14 | 4 | 14 |
| Lateral Ventricle* | | 7.99 | -8 | 2 | 4 |
| Left Amygdala | | 7.95 | -20 | -8 | -12 |
| Frontal Operculum Cortex | 1849 | 8.57 | -44 | 22 | 2 |
| Frontal Orbital Cortex | | 7.88 | -44 | 22 | -8 |
| | | 7.84 | -28 | 18 | -16 |
| | | 7.73 | -38 | 26 | -14 |
| Inferior Frontal Gyrus, pars opercularis | | 7.31 | -48 | 12 | 6 |
| | | 6.39 | -48 | 8 | 16 |
| Insular Cortex | | 6.38 | -28 | 26 | -2 |
| Inferior Frontal Gyrus, pars triangularis | | 6.21 | -44 | 32 | 12 |
| | | 5.65 | -46 | 34 | 0 |
| Frontal Pole | | 5.46 | -48 | 44 | 4 |
| Supramarginal Gyrus, posterior division | 527 | 7.72 | 66 | -44 | 20 |
| Supramarginal Gyrus, posterior division | | 6.49 | 50 | -42 | 14 |
| | | 6.45 | 52 | -44 | 22 |
| Middle Temporal Gyrus, posterior division | | 5.78 | 48 | -28 | -4 |
| Superior Temporal Gyrus, posterior division | | 5.73 | 48 | -34 | 2 |
| Precuneus Cortex | 351 | 7.52 | 2 | -58 | 32 |
| Supramarginal Gyrus, posterior division | 308 | 7.38 | -56 | -44 | 32 |
| Ventral Diencephalon* | 155 | 7.09 | -8 | -16 | -14 |

| | | | | | |
|---|---|---|---|---|---|
| Brain-Stem | | 6.75 | -2 | -26 | -2 |
| | | 5.29 | -6 | -28 | -16 |
| Frontal Orbital Cortex | 605 | 6.82 | 30 | 20 | -16 |
| | | 6.73 | 40 | 28 | -12 |
| Inferior Frontal Gyrus, pars triangularis | | 6.53 | 48 | 26 | 2 |
| | | 5.98 | 56 | 30 | 2 |
| Middle Temporal Gyrus, temporooccipital part | 294 | 6.75 | -56 | -58 | 8 |
| Right Hippocampus* | 104 | 6.39 | 30 | -10 | -12 |
| Right Amygdala | | 5.74 | 20 | -6 | -12 |
| Ventral Diencephalon* | | 5.37 | 12 | -6 | -10 |
| Right Amygdala | | 5.33 | 30 | -2 | -18 |
| Cingulate Gyrus, posterior division | 35 | 5.95 | -2 | -16 | 42 |
| Ventral Diencephalon* | 23 | 5.66 | 10 | -12 | -12 |
| Temporal Occipital Fusiform Cortex | 25 | 5.61 | -38 | -48 | -14 |
| Precentral Gyrus | 11 | 5.60 | 50 | 2 | 52 |
| Occipital Fusiform Gyrus | 167 | 5.58 | 30 | -88 | -10 |
| Occipital Pole | | 5.46 | 24 | -90 | -4 |
| Occipital Pole | | 5.31 | 28 | -92 | 4 |
| Occipital Fusiform Gyrus | 44 | 5.56 | -26 | -88 | -10 |
| Middle Frontal Gyrus | 21 | 5.31 | -30 | -2 | 52 |
| | 19 | 5.25 | -38 | 32 | 40 |
| Cingulate Gyrus, anterior division | | 5.19 | 0 | 22 | 20 |

| | | | | | |
|---|---|---|---|---|---|
| Angular Gyrus | 12 | 5.11 | 42 | -56 | 18 |
| Temporal Pole | 17 | 5.11 | 48 | 14 | -30 |
| Temporal Pole | | 4.84 | 40 | 16 | -30 |
| Middle Frontal Gyrus | 25 | 5.06 | -38 | 16 | 30 |
| Lateral Occipital Cortex, superior division | 28 | 5.06 | -40 | -64 | 20 |
| Lateral Occipital Cortex, inferior division | 8 | 4.87 | -24 | -90 | 4 |
| *Dishonest scenarios > Neutral scenarios* | | | | | |
| Superior Frontal Gyrus | 10997 | 11.91 | 0 | 46 | 38 |
| Middle Frontal Gyrus | | 10.88 | -38 | 16 | 44 |
| Superior Frontal Gyrus | | 10.80 | 0 | 54 | 32 |
| | | 10.49 | -8 | 26 | 58 |
| Frontal Pole | | 10.45 | 6 | 44 | 52 |
| Inferior Frontal Gyrus, pars triangularis | | 9.58 | -52 | 22 | 8 |
| Paracingulate Gyrus | | 9.25 | -6 | 20 | 48 |
| Middle Frontal Gyrus | | 9.24 | -40 | 4 | 50 |
| Frontal Pole | | 9.22 | -8 | 40 | 52 |
| | | 8.88 | -20 | 54 | 28 |
| Angular Gyrus | 3523 | 11.65 | -50 | -56 | 26 |
| Middle Temporal Gyrus, posterior division | | 9.83 | -56 | -24 | -8 |
| Angular Gyrus | | 9.45 | -52 | -58 | 38 |
| Temporal Pole | | 8.35 | -52 | 6 | -22 |

| | | | | | |
|---|---|---|---|---|---|
| Middle Temporal Gyrus, posterior division | | 7.63 | -52 | -38 | 0 |
| Middle Temporal Gyrus, anterior division | | 6.95 | -50 | -4 | -24 |
| Angular Gyrus | | 6.81 | -46 | -54 | 54 |
| Lateral Occipital Cortex, superior division | | 5.46 | -30 | -68 | 56 |
| Angular Gyrus | 1160 | 10.07 | 54 | -56 | 32 |
| | | 7.25 | 50 | -54 | 44 |
| Supramarginal Gyrus, posterior division | | 5.76 | 50 | -44 | 54 |
| Angular Gyrus | | 4.98 | 46 | -52 | 56 |
| Right Caudate | 1724 | 9.73 | 12 | 12 | 10 |
| Left Caudate | | 9.66 | -12 | 8 | 12 |
| Left Thalamus | | 8.25 | -8 | -14 | 12 |
| Right Thalamus | | 7.63 | 6 | -6 | 2 |
| | | 7.42 | 10 | -12 | 12 |
| Left Thalamus | | 7.17 | -4 | -8 | 4 |
| Ventral Diencephalon | | 7.05 | 6 | -10 | -10 |
| Left Caudate* | | 6.71 | -8 | 2 | 4 |
| Right Thalamus | | 6.62 | 4 | -26 | 2 |
| Brain-Stem | | 6.58 | -2 | -34 | -2 |
| Precuneous Cortex | | 9.21 | 0 | -60 | 32 |
| | | 8.69 | -2 | -56 | 40 |
| Cingulate Gyrus, posterior division | | 7.56 | 0 | -50 | 24 |
| Middle Temporal Gyrus, anterior division | 321 | 8.83 | 54 | 4 | -30 |

| | | | | | |
|---|---|---|---|---|---|
| Middle Temporal Gyrus, posterior division | | 5.56 | 60 | -8 | -26 |
| Frontal Orbital Cortex | 208 | 8.00 | 30 | 20 | -16 |
| | | 5.89 | 44 | 28 | -12 |
| Insular Cortex | | 4.95 | 30 | 10 | -6 |
| Cingulate Gyrus, posterior division | 109 | 7.97 | 0 | -20 | 40 |
| Middle Temporal Gyrus, posterior division | 396 | 7.89 | 66 | -20 | -10 |
| | | 6.69 | 58 | -30 | -6 |
| | | 6.32 | 54 | -22 | -10 |
| Occipital Pole | 559 | 6.64 | 26 | -94 | -4 |
| Lateral Occipital Cortex, inferior division | | 6.57 | 38 | -84 | -10 |
| Occipital Fusiform Gyrus | | 6.47 | 30 | -88 | -10 |
| Lateral Occipital Cortex, inferior division | | 6.02 | 32 | -88 | -2 |
| Occipital Pole | | 5.93 | 32 | -92 | 6 |
| Occipital Fusiform Gyrus | | 5.61 | 36 | -72 | -14 |
| Lateral Occipital Cortex, inferior division | | 4.94 | 46 | -80 | -2 |
| Left Putamen | 7 | 6.10 | -18 | 6 | 4 |
| Ventral Diencephalon* | 19 | 5.97 | -8 | -14 | -12 |
| Frontal Pole | 20 | 5.54 | -46 | 48 | 0 |
| Occipital Fusiform Gyrus | | 5.43 | -26 | -90 | -10 |
| Temporal Occipital Fusiform Cortex | 13 | 5.31 | 44 | -58 | -20 |
| Temporal Occipital Fusiform Cortex | 11 | 5.30 | -40 | -58 | -20 |

| | | | | | |
|---|---|---|---|---|---|
| Frontal Pole | 10 | 5.12 | 50 | 46 | -4 |
| Temporal Pole | 5 | 4.95 | 58 | 6 | -14 |

*location not included in the Harvard-Oxford atlas, and therefore given according to the SPM atlas

**Supplementary Table 2**. MNI coordinates for the peak activations (up to 10 peaks per cluster that are at least 8 mm apart, thresholded at 5 voxels) for the conjunction of moral judgement of disgusting, harmful and dishonest scenarios ($T$ (1, 245) = 4.77, Voxel-wise FWE corrected $p <$ .05). Region labels are according to the Harvard-Oxford Atlas.

| **Region Label** | **voxels** | **t-value** | **x** | **y** | **z** |
|---|---|---|---|---|---|
| Superior Frontal Gyrus | 2647 | 10.14 | -4 | 54 | 32 |
| Paracingulate Gyrus | | 8.70 | -6 | 52 | 18 |
| Superior Frontal Gyrus | | 8.01 | -6 | 20 | 56 |
| | | 7.19 | -4 | 46 | 42 |
| Frontal Pole | | 6.87 | -20 | 52 | 30 |
| Paracingulate Gyrus | | 6.72 | -6 | 34 | 36 |
| Superior Frontal Gyrus | | 6.25 | -6 | 28 | 44 |
| | | 6.10 | 6 | 20 | 58 |

| | | | | | |
|---|---|---|---|---|---|
| | | 6.06 | 10 | 18 | 66 |
| Frontal Pole | | 5.84 | -24 | 54 | 22 |
| Right Caudate | 871 | 8.73 | 12 | 12 | 6 |
| Left Caudate | | 8.42 | -12 | 10 | 8 |
| Right Caudate | | 8.22 | 14 | 4 | 14 |
| Left Thalamus | | 7.17 | -4 | -8 | 4 |
| Right Thalamus | | 7.10 | 6 | -6 | 4 |
| Left Caudate* | | 6.71 | -8 | 2 | 4 |
| Right Thalamus | | 5.75 | 2 | -14 | 6 |
| Inferior Frontal Gyrus. pars triangularis | 922 | 7.61 | -46 | 24 | 6 |
| Frontal Orbital Cortex | | 7.34 | -30 | 18 | -16 |
| Inferior Frontal Gyrus. pars opercularis | | 6.31 | -50 | 14 | 12 |
| Frontal Orbital Cortex | | 6.22 | -44 | 24 | -10 |
| Insular Cortex | | 6.18 | -28 | 24 | -2 |
| Inferior Frontal Gyrus. pars triangularis | | 5.11 | -50 | 34 | 0 |
| Precuneous Cortex | 334 | 7.52 | 2 | -58 | 32 |

| | | | | | |
|---|---|---|---|---|---|
| Frontal Orbital Cortex | 151 | 6.82 | 30 | 20 | -16 |
| | | 5.82 | 44 | 28 | -12 |
| | | 5.73 | 36 | 26 | -14 |
| Supramarginal Gyrus, posterior division | 146 | 6.57 | -58 | -48 | 32 |
| Left Putamen | 7 | 6.10 | -18 | 6 | 4 |
| Angular Gyrus | 125 | 6.06 | -56 | -58 | 14 |
| Left Thalamus | 57 | 5.99 | -4 | -24 | -4 |
| Right Thalamus | | 5.41 | 4 | -24 | -2 |
| Ventral Diencephalon* | 19 | 5.97 | -8 | -14 | -12 |
| Ventral Diencephalon* | 14 | 5.66 | 10 | -12 | -12 |
| Angular Gyrus | 9 | 5.60 | 64 | -48 | 22 |
| Occipital Fusiform Gyrus | 147 | 5.58 | 30 | -88 | -10 |
| Occipital Pole | | 5.46 | 24 | -90 | -4 |
| Cingulate Gyrus. anterior division | 21 | 5.57 | -2 | -14 | 40 |
| Inferior Frontal Gyrus. pars triangularis | 37 | 5.49 | 52 | 24 | 8 |

| | | | | | |
|---|---|---|---|---|---|
| Middle Temporal Gyrus. posterior division | 39 | 5.44 | 50 | -28 | -6 |
| Occipital Fusiform Gyrus | 15 | 5.43 | -26 | -90 | -10 |
| Temporal Pole | 13 | 5.11 | 48 | 14 | -30 |
| Middle Frontal Gyrus | | 5.07 | -34 | 0 | 58 |
| | 25 | 5.06 | -38 | 16 | 30 |
| Lateral Occipital Cortex. superior division | 12 | 4.92 | -46 | -66 | 20 |
| | | 4.81 | -38 | -62 | 20 |

*location not included in the Harvard-Oxford atlas, and therefore given according to the SPM atlas

**Supplementary Table 3**. MNI coordinates for the peak activations (up to 10 peaks per cluster that are at least 8 mm apart, thresholded at 5 voxels) for conjunctions of moral judgement of food-wasting and disgusting scenarios, food-wasting and harmful scenarios, as well as food-wasting and dishonest scenarios, each masked exclusively with the conjunction of moral judgement of disgusting, harmful and dishonest scenarios ($T$ (1, 245) = 4.77, Voxel-wise FWE corrected $p <$ .05). Region labels are according to the Harvard-Oxford Atlas.

| **Region Label** | **voxels** | **t-value** | **x** | **y** | **z** |
|---|---|---|---|---|---|

| | | | | | |
|---|---|---|---|---|---|
| *Food-wasting scenarios & Disgusting scenarios* | | | | | |
| Superior Frontal Gyrus | 2270 | 7.82 | 6 | 44 | 50 |
| Frontal Pole | | 7.78 | 10 | 50 | 44 |
| Superior Frontal Gyrus | | 7.73 | 18 | 34 | 54 |
| | | 7.68 | -18 | 34 | 54 |
| Frontal Pole | | 7.38 | -8 | 40 | 54 |
| Superior Frontal Gyrus | | 6.93 | 2 | 42 | 42 |
| Frontal Pole | | 6.86 | -14 | 46 | 44 |
| Middle Frontal Gyrus | | 6.75 | -36 | 18 | 48 |
| Superior Frontal Gyrus | | 6.71 | 12 | 30 | 58 |
| | | 6.65 | -12 | 24 | 64 |
| Lateral Occipital Cortex, superior division | 412 | 7.37 | 54 | -60 | 36 |
| | | 7.18 | 56 | -62 | 28 |

| | | | | | |
|---|---|---|---|---|---|
| Precuneous Cortex | 566 | 7.19 | -2 | -54 | 14 |
| | | 6.40 | -2 | -62 | 38 |
| | | 5.75 | -10 | -52 | 38 |
| | | 5.29 | -8 | -56 | 26 |
| Middle Frontal Gyrus | 113 | 6.90 | 40 | 22 | 52 |
| Inferior Frontal Gyrus, pars triangularis | 250 | 6.66 | -50 | 36 | 12 |
| Inferior Frontal Gyrus, pars opercularis | | 5.41 | -52 | 22 | 20 |
| Occipital Pole | 106 | 6.33 | 28 | -94 | -4 |
| Lateral Occipital Cortex, inferior division | | 5.30 | 36 | -88 | 0 |

| | | | | | |
|---|---|---|---|---|---|
| Occipital Fusiform Gyrus | | 5.02 | 22 | -88 | -4 |
| Occipital Pole | | 4.98 | 36 | -90 | 8 |
| Lateral Occipital Cortex, superior division | 504 | 6.22 | -42 | -70 | 40 |
| | | 6.22 | -56 | -62 | 24 |
| Angular Gyrus | | 5.56 | -44 | -58 | 34 |
| | | 5.42 | -50 | -58 | 44 |
| Lateral Occipital Cortex, superior division | | 5.26 | -50 | -70 | 32 |
| | | 5.24 | -34 | -74 | 48 |
| Left Caudate | 35 | 6.20 | -8 | 12 | 10 |

| | | | | | |
|---|---|---|---|---|---|
| | | 6.15 | -12 | 2 | 18 |
| Right Caudate | 26 | 6.09 | 8 | 12 | 10 |
| | | 5.42 | 12 | 6 | 18 |
| Paracingulate Gyrus | 40 | 5.66 | 0 | 48 | 24 |
| | | 5.24 | 8 | 48 | 18 |
| Left Amygdala | 26 | 5.55 | -24 | -6 | -16 |
| Right Thalamus | 24 | 5.50 | 10 | -10 | 10 |
| Left Thalamus | 26 | 5.49 | -8 | -16 | 12 |
| Lingual Gyrus | 21 | 5.44 | -4 | -86 | -6 |
| Left Thalamus | 7 | 5.37 | -8 | -10 | 0 |
| Left Putamen | | 5.33 | -16 | 8 | -2 |
| Cingulate Gyrus, anterior division | 14 | 5.20 | 0 | -14 | 34 |

| | | | | | |
|---|---|---|---|---|---|
| Frontal Pole | 10 | 5.18 | -20 | 58 | 24 |
| Lateral Occipital Cortex, inferior division | 5 | 5.14 | 40 | -82 | -10 |
| Cingulate Gyrus, posterior division | 20 | 5.07 | 0 | -34 | 30 |
| Right Amygdala | 5 | 5.04 | 22 | -4 | -18 |
| Frontal Orbital Cortex | | 4.80 | 42 | 34 | -14 |
| *Food-wasting & Harmful scenarios* | | | | | |
| Inferior Frontal Gyrus, pars triangularis | 164 | 6.21 | -44 | 32 | 12 |
| Frontal Pole | | 5.57 | -48 | 38 | 6 |
| Superior Frontal Gyrus | 14 | 6.01 | 12 | 20 | 60 |
| | 24 | 5.65 | -12 | 16 | 60 |
| | | 5.07 | -12 | 10 | 66 |
| Left Amygdala | 13 | 5.38 | -22 | -6 | -14 |

| | | | | | |
|---|---|---|---|---|---|
| Left Putamen | 7 | 5.33 | -16 | 8 | -2 |
| *Food-wasting & Dishonest scenarios* | | | | | |
| Superior Frontal Gyrus | 2588 | 8.47 | -14 | 34 | 56 |
| | | 8.06 | 12 | 36 | 52 |
| | | 8.00 | -16 | 24 | 62 |
| | | 7.88 | -12 | 26 | 54 |
| | | 7.82 | 6 | 44 | 50 |
| Frontal Pole | | 7.77 | 8 | 50 | 44 |
| | | 7.77 | -10 | 40 | 52 |
| Middle Frontal Gyrus | | 7.30 | 38 | 22 | 50 |
| Superior Frontal Gyrus | | 7.21 | 16 | 22 | 60 |
| | | 6.93 | 2 | 42 | 42 |
| Lateral Occipital Cortex, superior division | 469 | 7.37 | 54 | -60 | 36 |

| | | | | | |
|---|---|---|---|---|---|
| | | 7.18 | 56 | -62 | 28 |
| | | 5.64 | 50 | -58 | 48 |
| Precuneous Cortex | 376 | 6.40 | -2 | -62 | 38 |
| Cingulate Gyrus, posterior division | | 5.94 | -2 | -52 | 20 |
| Precuneous Cortex | | 5.75 | -10 | -52 | 38 |
| Cingulate Gyrus, posterior division | | 5.22 | -10 | -52 | 28 |
| Occipital Pole | 100 | 6.33 | 28 | -94 | -4 |
| Lateral Occipital Cortex, inferior division | | 5.15 | 34 | -88 | 2 |
| Occipital Fusiform Gyrus | | 5.02 | 22 | -88 | -4 |

| | | | | | |
|---|---|---|---|---|---|
| Lateral Occipital Cortex, superior division | 414 | 6.22 | -56 | -62 | 24 |
| | | 5.56 | -42 | -68 | 40 |
| Angular Gyrus | | 5.56 | -44 | -58 | 34 |
| | | 5.53 | -50 | -58 | 44 |
| Lateral Occipital Cortex, superior division | | 5.26 | -50 | -70 | 32 |
| Left Caudate | 35 | 6.20 | -8 | 12 | 10 |
| | | 6.15 | -12 | 2 | 18 |
| Right Caudate | 22 | 6.09 | 8 | 12 | 10 |
| | | 5.42 | 12 | 6 | 18 |
| Paracingulate Gyrus | 40 | 5.66 | 0 | 48 | 24 |
| | | 5.24 | 8 | 48 | 18 |

| | | | | | |
|---|---|---|---|---|---|
| Inferior Frontal Gyrus, pars triangularis | 69 | 5.55 | -52 | 30 | 16 |
| Inferior Frontal Gyrus, pars opercularis | | 5.41 | -52 | 22 | 20 |
| Right Thalamus | 24 | 5.50 | 10 | -10 | 10 |
| Left Thalamus | 26 | 5.49 | -8 | -16 | 12 |
| Middle Temporal Gyrus, posterior division | 17 | 5.40 | 56 | -8 | -28 |
| Frontal Pole | 10 | 5.18 | -20 | 58 | 24 |
| Lateral Occipital Cortex, inferior division | 5 | 5.14 | 40 | -82 | -10 |
| Cingulate Gyrus, posterior division | 7 | 5.08 | -2 | -16 | 36 |

*location not included in the Harvard-Oxford atlas, and therefore given according to the SPM atlas

**Supplementary Table 4.** MNI coordinates for the peak activations (up to 10 peaks per cluster that are at least 8 mm apart, thresholded at 5 voxels) for the contrast of food-wasting against disgusting, harmful and dishonest scenarios pooled together ($T$ (1, 245) = 4.77, Voxel-wise FWE corrected $p < .05$). Region labels are according to the Harvard-Oxford Atlas.

| **Region Label** | **voxels** | **t-value** | **x** | **y** | **z** |
|---|---|---|---|---|---|
| Parahippocampal Gyrus, posterior division | 76 | 7.03 | -28 | -32 | -20 |
| Precuneous Cortex | 106 | 7.01 | -10 | -58 | 16 |
| Insular Cortex | 21 | 5.63 | -38 | -6 | 6 |
| Frontal Orbital Cortex | 5 | 5.35 | 22 | 34 | -14 |
| | 6 | 5.27 | -28 | 34 | -12 |
| Middle Frontal Gyrus | 15 | 5.09 | 34 | 22 | 52 |
| Frontal Pole | 8 | 4.99 | 44 | 38 | 20 |

### Stimuli – food-wasting, moral transgressions and neutral scenarios (translated into English)

#### I. Food-wasting scenarios

1. Olivia visits her parents in the countryside. At the end of the visit, she receives a huge bag of goods – frozen dumplings, meat, and jars with preserves. Olivia does not like being

called a country bumpkin in the city by her flatmates, so just after leaving the train, she throws the bag into a mixed waste container.

2. Robert is going grocery shopping for the family to buy groceries for the whole week. This time, he forgot to take the shopping list from his wife. After returning home, it turns out that the man bought too much food. In order to avoid an argument with a nervous spouse, Robert quickly goes to the trash and throws away the unnecessarily purchased groceries.
3. Harry notes that he can benefit from a supermarket promotion in which two loaves of a certain bread are cheaper than one loaf. Harry needs only one loaf. However, he buys two at a lower price, and throws the one he does not need straight into the trash outside of the store.
4. Amelia is really bored because there are no other children in the yard that day. The girl comes up with the idea of buying a dozen loaves of bread and kicking them like balls. Amelia enjoys herself throughout the afternoon by destroying the loaves she bought. After playing, she throws the leftover bread into the trash.
5. Emily is on a diet. She asks her boyfriend to do the shopping for her for the next month at the wholesale market. When the boy returns with the shopping, the girl realizes that he bought 8 liters of milk which has a high fat content. Emily sends her boyfriend for skim milk and then pours all of the high fat milk into the sink.
6. Jack buys a few dozen kilograms of pork from the butcher to make home-made sausage. The man uses only part of the purchased goods, because during work he realizes that it takes too much time and he does not have enough time to rest. Jack throws the unused pork into the trash.

7. Alice is the owner of a dairy which produces large amounts of milk, cheeses, yoghurts, and other dairy products every year. However, it often happens that the company produces more dairy products than it is able to sell. Alice orders excess milk to be poured into the drains and cheese and other products to be dumped into trash.
8. George is on a business trip in India and leaves the hotel for dinner at street vendors. The dishes are so tasty and cheap that he buys a few bags full of food to try several different dishes. After a while, he is already full, so he throws two full bags of food into the trash.
9. Grace is a supermarket manager. She orders her employees to throw away all fruits and vegetables that were not sold the day before. Every day, huge amounts of food are taken to the trash. Grace is satisfied because only fresh goods remain on the shelves of the vegetable department.
10. Mark goes out to dinner with a girl he met at work. He wants to impress her, so when ordering food, he decides to get a large number of expensive dishes. When the waiter starts bringing the food, Mark says that he is not hungry and orders the waiter to throw all the food into the trash.
11. Jacob buys fresh bread in the bakery. After returning home, the boy is terribly hungry, so he makes two plates of sandwiches with cheese, ham, and tomato. After taking the first bite, he is repelled by the taste of cumin which is in the bread. As a result, he throws all of the sandwiches into the trash.
12. Sophie organizes a scientific conference for which she orders snacks from a catering company – sandwiches, pastries and cakes. There is a lot of food left and there is no refrigerator on the premises. Sophie thinks about giving the snacks to the students, but she is in a hurry to get home, so she throws everything that’s left into the trash.

**II. Disgusting scenarios**

1. Jessica is going to a party with her brother. The boy drinks too much alcohol and falls asleep on the couch. When no one is in the room, Jessica starts kissing him and grabbing him by his crotch through the trousers. This lasts only a few seconds and the brother is not aware of anything.
2. Noah decides to spend the upcoming weekend in a local public house. During sexual intercourse with a selected prostitute, it turns out that it is his younger sister who moved out of the house some time ago. Noah pretends that he has not noticed anything and continues to have sex.
3. Ruby is a young girl who has a way with men. One day, she and her friend make a bet that she will persuade each man encountered in the nearby park to have oral sex with her. When Ruby’s brother is passing through the park, in order not to lose a bet, the girl has oral sex with him.
4. Poppy decides to visit her sister, whom she has not seen in twenty years. The sister is not at home, but a young man invites Poppy in to wait. Poppy seduces the boy and they have sex. Poppy is not surprised when it turns out that the young lover is her nephew.
5. Freya is professionally involved in dance. However, it is hard for her to make a living with money she earns at shows, competitions, and festivals. Freya decides to try her luck as an erotic dancer. She performs as a stripper at her brother’s bachelor party, where she kisses him passionately on the table while almost naked.

6. Sara, after graduation, lives with her older brother in one apartment in a big block of flats. When no one is around, sometimes they happen to touch their genitals passionately and kiss each other. This behavior is not a problem for either of them.
7. Benjamin visits his 67-year-old grandmother. When the elderly woman is cooking, her grandson approaches her from the back and begins to kiss her passionately. They both feel arousal and start rubbing against each other until they reach an orgasm. Neither of them sees any problem in this situation.
8. Theo loves sailing with his brother Bobby. One night, when the brothers are on a yacht together, they engage in anal intercourse. They both feel pleasure thanks to this, and the memory of the event makes them feel closer. What happened never brings any negative consequences to them.
9. Phoebe is going with her friends to a party in a club where she meets a handsome man. After having a few drinks, the girl and her companion sit on the couch together and kiss for the rest of the party. Phoebe is particularly attracted to him due to the fact that he might be her close relative.
10. Charlie is a young man. He has just graduated from university and is about to begin his first serious job. He decides that the time has come to finally settle down and start a family. Shortly after this decision, he marries his girlfriend who happens to be his half-sister.
11. Mason is at a family meeting in a mountain hotel. At some point, he meets an attractive woman he has never seen before. The woman turns out to be his close cousin who lives on another continent. They have a nice conversation and after a few drinks they move to the bedroom, where they have sex.

12. Thomas was given up for adoption shortly after birth and he has never met his biological parents. One day, during a choir rehearsal, he meets an attractive older woman. He invites her to dinner, which ends in sex. During sex, Thomas recognizes that the woman is his biological mother, whose photo he keeps in his wallet.

**III. Harmful scenarios**

13. Peter is returning from a birthday party through the park. He notices that there is a person lying in the bushes. He approaches and sees that it is a drunk homeless person who has been seriously beaten. Peter kicks him with all his strength in the stomach and starts walking away towards his home with no rush.
14. Layla likes walking with her little dog in the park. However, she hates it when other dogs run free and want to sniff her pet. Therefore, every now and then Layla scatters sausage with poison in the park, hoping that strange dogs will eat it and this way she will get rid of the problem.
15. Natalie is an owner of a purebred dog. However, nobody in the family has time to take care of him and, according to Natalie, the expenses for food and veteranarian are too high. Therefore, the woman takes the dog into the forest and lets him free, hoping that the animal will find a new home.
16. Archie is bored after yet another day on holiday at his grandparents' house in the countryside. He decides to have some fun.  He goes outside and comes up with the idea to pick up stones and throw them at the cows grazing on a nearby pasture. He is having great fun while doing this.

17. Margaret is going for a parent - teacher meeting at school. She hears that her 8-year-old son is the worst in the class in math and he does not do his best during class. After returning home, she enters her child's room and slaps him with her open hand in his face, punishing him for his poor grades.
18. Edward works in an industrial slaughterhouse. His main task is bringing the pigs into the room in which they will be stunned by a pneumatic gun. Edward does not like waiting until the pigs enter the room willingly, so he often kicks them, burns them with a cigarette, and hurries them with a wooden stick with a sharp spike at the end.
19. Maria is standing in a long queue at the supermarket. She is very bored and does not have a phone to play a game. She surreptitiously turns over the shopping basket of an elderly woman who is standing in front of her and when the woman starts collecting her groceries, Maria laughs at her own joke.
20. Finley is a teenager who is extremely popular at school. One day he notices that one of his classmates comes to school in a dirty, cheap sweatshirt. Finley makes fun of him and tells everyone that this colleague is poor and dirty and leaves in a pigsty.
21. Annie organizes a surprise birthday party for her younger brother. Annie knows that her brother is terribly afraid of clowns and darkness. Nevertheless, she asks all guests to come in clown costumes. At the beginning of the party she plans to turn off the lights and play music from a horror movie.
22. Elisabeth lives at home with her daughter and grandchildren. One of her grandchildren is running around the house and making a terrible noise. Elisabeth approaches him and whispers to him, looking into his eyes, that if the child does not calm down, then she will come to his room at night and will strangle him with a pillow.

23. Richard is a manager of a company. One day, he decides to play a joke on his employee who has just taken a mortgage. He approaches her and announces that she is fired and that she is to pack immediately. The employee starts crying. Richard cannot stop from laughing out loud.
24. Finn returns home from work in a bad mood and is waiting for dinner that his wife is making. After eating a few bites, he yells at the woman that the food is disgusting and that his wife cannot cook. Then, the man leaves the house, slamming the door, to eat a kebab with fries.

**IV. Dishonest scenarios**

25. Alice orders goods for her company worth $3530. When the invoice comes, the amount to be paid is $3503, and not $3530. Alice knows that the supplier of goods made a mistake but decides that she will not tell anyone about this mistake.
26. Daisy is invited to dinner at her grandparents' house. However, she is not willing to visit them. So she calls her grandmother and tells her that she is not feeling well and that she has a fever. Grandmother, very worried, tells Daisy to stay at home and to take the right medication.
27. Mia borrows a car from her boyfriend to go shopping. While backing up, she accidentally drives into a post and scratches the side of the car. She knows that her boyfriend will be furious when he finds out what happened, so she tells him that the scratch was already on the car when she left the store.
28. Scarlett is a player in a very well-known professional basketball team. During the final of the most important match of the season, Scarlett realizes that the team needs some help,

because the match may end in a loss. At some point, the woman pretends that she is fouled. Thanks to this, her team gains additional points and wins the championship of the country.

29. Chris is shopping for a lot of groceries. When paying, he does not notice a smartphone at the bottom of the shipping cart. Chris realizes that he did not pay for the product only when repacking the groceries from the cart into the car. The man decides he will not go back to the store to pay for the new phone.
30. Samuel is a young student living in a big city. He uses public transport every day and, even though he does not run out of money, he never buys tickets. When he encounters a ticket controller, he gets off the bus and runs immediately. Samuel has never received a ticket.
31. Ella works as a florist. One day, she gets an order for a bouquet of 300 roses. Ella consciously makes a bouquet composed only of 250 roses and issues a receipt for the customer that amounts to 300 roses. Nobody else notices that the bouquet is smaller than it should be.
32. Mary works in the office. When her supervisor asks her how many hours she has worked in previous weeks, Mary knows that her boss does not control her working hours and will not notice if she provides an overstated number. Mary replies that she worked much more than she actually did. She gets a raise for the supposedly hard work.
33. Logan works in a large office. He does not want to take lunch with him to work. Instead, he eats other employees' food. So that no one knows, he always takes very small amounts of other people's meals. Colleagues do not notice that Logan steals their food.
34. Noah gets $300 from his grandmother. His grandmother asks him to share the money with two siblings – a younger brother and a twin sister. Noah keeps $200 for himself and gives $50 to each sibling. He tells his grandmother that he divided the money equally.

35. Andrew is selling an old car. He knows that if he tells the client that his car was in a collision, the price of the car will drop. Andrew manages to sell the car to a young boy who does not find out that the car was fixed. Thanks to this, Andrew gets more money for selling the car.
36. Mark is a judge with many years of experience. One time, he got a case in which a woman who was his youthful love was the accused. Mark spends the evening reminiscing about the old days with a glass of wine. On the day of the trial, despite the obvious fault of the woman, he acquits his past lover.

**V. Neutral scenarios**

37. Jack is going to his friend's birthday party. He wants to surprise his friend, so he prepares a fantastic cake with fresh strawberries and a lot of cream. While cooking, his favourite program is playing on the radio and when kneading the dough, the boy is singing loudly and stamping his foot to the rhythm of the music. It does not seem to bother anyone.
38. Annie is organizing a school collection for a shelter for homeless animals. For this purpose, she draws beautiful portraits of dogs and cats. The next day she goes from class to class collecting money for food for the animals. She gives one of the portraits to each child who throws a coin into the piggy bank.
39. William is planning to stay home for the holidays to spend time with his childhood friends. William's father had a car accident, as a result of which he suffered complex fractures in both legs. William will be happy to help his father in his daily duties while he gets better.
40. Abigail wants to adopt a rescue dog. Browsing the websites of the shelters, she finds exactly the dog she was looking for. The shelter, where the animal is staying, is located 50

kilometres from Abigail’s place of residence. After giving thought to it, the woman borrows a car from a friend and drives to meet the dog she found.

41. Ezra needs to transport a large box from a tool shop, but the load does not fit into his car. So he asks Nick if he could borrow his delivery truck. Nick agrees. So Ezra transports the box, then gives the car back to Nick with a full tank of fuel and without any damage.
42. Steve sings beautifully and his tutor persuades him to enrol in a music school. Steve tells his parents about this. They agree to enrol their son in singing lessons. Steve is very happy and gratefully prepares a song for his teacher.
43. Benjamin is a programmer. He sits in front of a computer all day which makes him very tired. Therefore, he decides that he will spend the evening away from the screens. He leaves all the electronic equipment at home, including the phone, and goes to kung-fu training and then to meet his friends.
44. Maryam notices that her dog is strangely dull and has a dry nose. So she calls the veteranarian and makes an appointment. It turns out that the dog is sick and must get an antibiotic. Over the following five days, Maryam gives her pet a pill with an antibiotic in a piece of ham.
45. Lewis is very concerned about the fate of animals bred for meat. After reading a report about industrial breeding of chickens, he decides to limit his consumption of poultry and eggs. Over the following days, he keeps his promise to himself and eats only vegan meals. He is very pleased.
46. Sebastian comes to the conclusion that he feels bad about the fact that he has not practiced sports for some time. He recalls that as a child he liked playing football and persuades his

friends to participate in regular matches in a nearby sports hall. His friends like his idea, so Sebastian books the hall.

47. Clara has always wanted to make tinctures. When autumn comes, she buys a kilogram of chokeberry, adds sugar, and pours alcohol over the mixture. After three months, when the time comes to open the bottle in which the chokeberry tincture was maturing, she discovers that her first home-made tincture is quite tasty.
48. Heidi is an teacher and gives her students the date of an exam. Two students cannot attend the exam on the specific date and ask her for the opportunity to take the test on another day. Heidi looks at her schedule and then offers her students a different date. They agree on the new date.
49. Barbara is a single older woman. On vacation in a seaside retreat, she meets a very interesting single man her age. They get along very well and Barbara is considering inviting him to visit her in her town. However, she comes to the conclusion that she must first get to know him better.
50. Katie is finishing her bachelor’s studies this year and she does not feel ready to start a professional career. She enrols for master’s degree studies, which seem very interesting to her and can help her finding an interesting job in the future. To make a living while studying, Katie works at night as a security guard at a construction site.
51. Molly would like to get into her dream university, so she studies hard for her final exams. She does not go to parties and a few months before the secondary school exam, she quit all of the extracurricular activities she attended. The time she gains in this way, she spends on additional courses preparing for the final exams.

52. Mila would like to take part in a film festival. However, the festival pass is very expensive. Mila learns that she can get a pass in exchange for volunteering during the festival, which involves providing service at a film store. This type of work fits Mila, so she volunteers.
53. Darcie works as a downhill skiing instructor in winter. To prepare herself physically for the season, she begins to train her thigh and calf muscles and to train her balance in September. In addition, she goes running in the park twice a week to feel better once she is on the slopes.
54. Amber makes a New Year's resolution to eat healthier. For this reason, before going to work, she prepares a salad to take with her to the university. At lunchtime, the woman eats what she prepared and enjoys the meal. At the end of the day, she is satisfied with the decision she made.
55. William is a drummer in a rock band. He usually practices the drums in a rented rehearsal room. He would also like to play at home, but  does not want to disturb his neighbours. So the man decides to buy digital drum set, which is quiet, because you can connect headphones to it.
56. Hannah has been thinking about buying a new coat for some time. When sale time comes, the girl finally finds time to go to the store and see the clothes being offered at discount. The coat she likes most is on sale. Hannah buys it and is happy that she saved money.
57. Daniel is getting married in August. Together with his fiancée, they book a date in the office and a venue for the wedding. Although it is only January, they send invitations to the relatives for the event. Daniel is personally asking that the invitees inform him one month before the wedding whether they will attend..

58. Ryan is a student at a technical university. Recently, he has been learning mathematics, chemistry, and physics every day because he is preparing for exams. After another day spent learning, in order to regain strength, he goes for a long walk. After returning home, he drinks bitter cocoa and takes a half-hour nap.
59. Alexander is going to an exquisite French restaurant for the first time in his life. On the menu he sees vichyssoise (cold potato soup). He does not know what vichyssoise is, but he orders the dish and eats it. He does not really like the soup, but he is actually very happy that he tried it.
60. Jasmine buys a beautiful fern in a shop. After returning home, she notices that the plant has a pot that is too small. She finds appropriate information online about the proper replanting of potted plants. She then goes to the store for a larger pot and replants her new fern.
61. Alice likes going for long walks in the park on Sunday morning with her friend Bonnie. This Sunday, Bonnie is visiting her family in a different city. Alice decides to go for a walk without Bonnie. The walk is very pleasant, although Alice misses talking to her friend.
62. Emma is leaving for three weeks for a therapeutic retreat. She does not want her potted plants to wither during this time, so she asks her neighbour, Ms. Beatrice, to water her plants once every three days. Ms. Beatrice promises to take care of the flowers and Emma leaves her the key to her apartment.
63. Nathan is preparing for his driver’s licence test. He buys additional driving lessons with the instructor and asks him for help in improving some manoeuvres which are still challenging to him. After two hours of driving, Nathan is happy and feels that he is doing much better with the manoeuvres he practiced.

64. Bobby hears on the radio that he could win several million dollars in a lottery. Although he usually does not do such things, he buys three tickets and marks random numbers. When the results of the lottery are announced in the evening, he tracks them with interest and does not worry that he did not get them right.
65. Nancy has worked a lot for the last few months and thinks about taking a vacation. She looks through offers of travel agencies and finds several interesting offers. The next day, the woman speaks with her boss, who encourages her to take time off and rest. Nancy buys one of the trips and begins planning the details of the journey.
66. Dexter has been planning to buy his own flat for some time. One day, he finds a property that meets his expectations and is reasonably priced. Dexter already has half of the money needed to buy a flat, so he makes an appointment with a bank advisor to learn about mortgage offers.
67. Robyn is meeting her friend Simon for coffee on Wednesday at noon. On Wednesday morning, Robyn does not feel well. The only available appointment to see a doctor is at noon. Robyn calls Simon, who agrees to postpone the meeting in the café until the next week.
68. Thea is planning to buy a used car. When she finds an interesting car, she asks the owner to go to a pre-sale service with her. The owner agrees and the car is checked by a car mechanic. When it turns out that everything is alright, Thea buys the car at the previously agreed price.
69. Ollie always comes back home from class by bus number 117. One day, as frequently occurs, the vehicle gets stuck in traffic. After returning home, Ollie, irritated by travelling

in a crowded bus, checks online whether it is possible to get home from the university using a tram.

70. Elliot spends his Saturday mornings cleaning his apartment. While vacuuming the floor in the bedroom, he notices that the light bulb in the lamp by his bed is not working. The man finishes vacuuming, then puts on his jacket and shoes, takes his wallet, and goes to a nearby store to buy a new light bulb.
71. Monica is driving to work. The cars in the lane where she is driving are moving forward slowly compared to cars in the next lane. Monica is already late for work, so after carefully looking in the mirror and turning on the turn-signal, she changes lanes to the faster one.
72. Megan did not manage to buy a tram ticket in the machine at the stop. After entering the tram, she goes to the machine located in the front of the vehicle and buys the appropriate ticket. To avoid losing her balance, she waits for the tram to finish turning and then validates the ticket she purchased.
73. Finley comes to the conclusion that the shoes he bought recently are one size too small. So he does not unpack them from the box and does not take the labels off. Thanks to the fact that he kept the receipt, he can exchange the shoes. So the man returns to the store and exchanges the shoes for another pair having the correct size.
74. Tyler notes that his eyes are sore and he gets a headache while reading. So he goes to have his eyes checked. The checkup shows that he should wear reading glasses. So he buys the frames in which he looks the best. When he starts reading, he puts the glasses on, according to the recommendations of the ophthalmologist.
75. Leon lives in the city center and cannot fall asleep because of the noise outside. He decides to close the window in the bedroom so that there is less noise. He wraps himself in a

dressing gown and gently closes the window, then climbs under the covers and falls asleep without any problems.

76. Matthew decides to bake cookies. So he goes to the store to buy the missing ingredients. After returning home, the man starts kneading the dough. He suddenly remembers that he forgot about the flour. After searching the cupboards, Matthew finds a whole bag of flour in one of them.
